# Going beyond labels: Semi-supervised learning for enzyme kinetic parameter prediction

**DOI:** 10.64898/2026.10.15.759462

**Authors:** Ali Malli, Denys Vasyutyn, Anuj Majumder, Jin Ryoun Kim

## Abstract

Enzyme kinetic parameters, such as the turnover number (*k*_cat_), underpin enzyme engineering and metabolic modeling. Machine learning models can predict these parameters from sequence and substrate representations, but their accuracy is limited by the scarcity and narrow coverage of experimentally labeled data, particularly for out-of-distribution enzymes. Existing strategies to exploit unlabeled data rely on homologous sequences, pseudo-labels or synthetic perturbations of labeled samples, limiting their potential to fully leverage unlabeled data for model improvement. Here, we show that regularization-based semi-supervised learning improves model performance by leveraging biologically meaningful unlabeled samples alongside labeled data during task-specific training, without assigning pseudo-labels to the unlabeled samples. Specifically, enzyme-substrate pairs lacking *k*_cat_, but characterized for other kinetic parameters, can serve as unlabeled data that improve *k*_cat_ prediction. Using the clustered CataPro dataset at 40% sequence identity as a stringent benchmark, neural consistency regularization and graph-Laplacian regularization each performed better when unlabeled records were added, beyond the gains from regularization on labeled data alone. The benefit required sufficient labeled data, saturated at a few thousand unlabeled records for the neural model, and depended on the density of the unlabeled data in the feature space rather than on their proximity to labeled data. Gains extended to wild-type and mutant enzymes and to a local low-data adenylate kinase landscape. These findings show that incomplete kinetic records specifically, and unlabeled data points generally, are a valuable resource for enzyme kinetics prediction.

## 1. Introduction

Kinetic parameters provide a quantitative description of an enzyme’s catalytic function, making them central to enzyme engineering, metabolic modeling, and enzyme mining. Nevertheless, the catalytic turnover number, *k*_cat_, and the Michaelis constant, *K*_m_, are available for only a small fraction of known enzyme-substrate pairs because their experimental determination is laborious and difficult to scale^1^. Although advances in laboratory automation and microfluidics have greatly increased the throughput of enzyme screening^2,3^, large-scale determination of kinetic parameters across broad sequence and substrate space remains considerably more challenging. Recent machine learning (ML) approaches have sought to address this limitation by predicting kinetic parameters from enzyme sequences and/or structures together with substrate representations^1,4^. Models such as UniKP^5^, CatPred^6^, and CataPro^7^ have demonstrated that pretrained protein and molecular representations can support this prediction task, even for enzyme sequences that are out-of-distribution relative to the training data^6,7^. Despite these advances, the performance of such models remains fundamentally constrained by the scarcity and limited coverage of experimentally measured kinetic labels.

Protein sequence data continue to accumulate far more rapidly than experimentally characterized functional data, with UniProt^8^ containing nearly 150 million protein sequence entries while experimentally measured enzyme kinetic parameters available in databases such as BRENDA^9^ remain orders of magnitude more limited^1^. Unlabeled data can provide one means of reducing this dependence on experimentally measured labels. Several strategies for incorporating such data have emerged in protein engineering. These methods differ in how they extract information from unlabeled data. Self-supervised learning uses large collections of natural protein sequences without functional labels to learn biologically informative representations that can subsequently be used in a supervised prediction task^10–14^. In contrast, semi-supervised learning (SSL) incorporates both labeled and unlabeled data into learning the task at hand, for example by assigning pseudo-labels to unlabeled data^15^ or by introducing a term pertaining to unlabeled data into the training objective^16^. Alternatively, synthetic data augmentation generates additional observations from existing labeled data rather than introducing independent unlabeled samples^17^.

An early example of the self-supervised learning strategy is the low-*N* protein-engineering framework developed by Biswas et al. (2021). UniRep^14^ was initially pretrained on millions of natural protein sequences, and subsequently fine-tuned on tens of thousands of unlabeled sequences related to *Aequorea victoria* green fluorescent protein (avGFP). A supervised regression model was subsequently trained to predict protein fitness using as few as 24 experimentally characterized variants^11^. For avGFP, the model enabled the identification of new variants with fluorescence exceeding that of the wild type and outperformed models that did not incorporate the local unlabeled sequence landscape^11^. Importantly, the unlabeled sequences were used to adapt the pretrained protein representation before supervised learning and did not participate directly in learning the fitness target. Thus, this strategy leverages unlabeled sequences primarily to specialize a pretrained representation to a target protein and its local evolutionary neighborhood, rather than incorporating unlabeled samples directly into prediction across a broad protein landscape.

Meanwhile, SSL has been explored for protein property prediction in different forms. ProteinNPT introduced a pseudo-generative transformer architecture that jointly models protein sequences and available experimental properties to improve prediction across label-scarce protein engineering tasks^18^. Specifically, this model has been applied to predict *k*_cat_ and *K*_m_ in adenylate kinases, outperforming a lightweight supervised regressor trained on the available labeled data^2^. While ProteinNPT benefits from pretrained embeddings on large-scale unlabeled protein sequences, it does not directly incorporate additional sequences lacking target labels into model training. Instead, it trains on labeled sequences while masking known labels and amino acids for target prediction and sequence reconstruction. Another class of SSL entails using wrapper methods where a supervised model first predicts the target for unlabeled samples, and high-confidence predictions are assigned as pseudo-labels that are incorporated into subsequent training rounds^15,19^. Olivares-Gil et al. (2025) evaluated Co-Training and Tri-Training regression^20^ using homologous protein sequences as unlabeled examples across 19 protein-fitness datasets. These wrapper methods provided less consistent improvements than unsupervised and self-supervised preprocessing approaches tested in the study^21^, attaining a Spearman’s correlation coefficient (*ρ*) comparable to the supervised baseline at best.

A separate line of work has addressed limited kinetic data through synthetic augmentation. O^2^DENet generates perturbed versions of labeled enzyme-substrate pairs through enzyme sequence and molecular graph masking. Then, it encourages agreement between the representations of the original and augmented samples^22^. This invariance objective improved enzyme kinetic prediction under distribution shift^22^, but the augmented samples remain synthetic derivatives of enzyme-substrate pairs for which kinetic measurements are already available. More recently, AUKAT used a conditional variational autoencoder to generate additional biochemical instances directly in the embedding space. The generated instances were assigned estimated *k*_cat_ values from neighboring labeled examples and filtered according to their agreement with an independent random forest regression model before being incorporated into the training set^23^. Although both approaches expand the training set, the additional training information is ultimately derived from the existing labeled kinetic dataset rather than from independently documented enzyme-substrate pairs whose target kinetic value is unknown. Hence, they expand the local neighborhood around labeled data, yet do not introduce new biochemical regions in the feature space.

Here, we investigate whether unlabeled enzyme-substrate data can improve enzyme kinetic prediction for observations with low similarity to the training data, which remains a highly challenging task. We evaluate this approach using two SSL mechanisms. First, we use neural consistency regularization, in which two independently perturbed views of the same enzyme-substrate representation are encouraged to produce similar predictions, allowing both labeled and unlabeled observations to influence model training without requiring an estimated kinetic target for the unlabeled data. Second, we evaluate a graph-based formulation in which enzyme-substrate observations are represented as nodes in a nearest-neighbor graph that leverages Laplacian regularization to encourage strongly connected observations to have similar inferred kinetic values. In this setting, unlabeled nodes can influence prediction through the geometry and connectivity of the combined labeled-unlabeled feature space while target fidelity is imposed only on labeled nodes. By testing these mechanistically distinct approaches, we ask whether any benefit from the unlabeled records reflects a general contribution of unlabeled data rather than an effect specific to a sole method. We distinguish the impact of regularization from the additional contribution of these unlabeled records by comparing a supervised baseline, a regularized model using labeled observations only, and a combined model incorporating both labeled and unlabeled observations under sequence-clustered cross-validation and repeated random seeds. To avoid introducing noise by arbitrarily pairing unlabeled enzyme sequences with substrates, we leveraged biologically grounded enzyme-substrate pairs for which the target kinetic parameter is missing, without assigning estimated kinetic labels. Because consistency and graph-based regularization can, in principle, be incorporated into a range of predictive models, the utility of unlabeled kinetic records may extend beyond the specific model architectures evaluated here.

## 2. Results

We evaluate the SSL formulations for enzyme kinetic parameter prediction on sequences that are distant from the training data. For this evaluation, we used the *k*_cat_ module in CataPro, which represents enzyme-substrate pairs using pretrained ProtT5 protein embeddings, MolT5 molecular embeddings, and MACCS fingerprints and provides ten predefined splits constructed under a 40% enzyme sequence identity threshold. The CataPro dataset comprises 13,211 unique enzyme sequences and 3,302 unique substrates spanning reactions across diverse EC groups. Although large sequence repositories such as UniProt provide a vast pool of uncharacterized protein sequences for SSL, many may not be readily expressible or active toward the substrates represented in CataPro. To identify more biologically meaningful unlabeled data, we selected enzyme-substrate pairs from CataPro that had been characterized for other kinetic parameters but lacked the target kinetic value. Enzyme-substrate records with experimentally measured *k*_cat_ values formed the labeled set whereas experimentally characterized pairs with a reported *K*_m_ but no corresponding *k*_cat_ formed the unlabeled set (Figure 1, a, b). Rather than introducing a new representation or model architecture, we retain the feature space and a simplified evaluation framework from CataPro to isolate the contribution of semi-supervised training (Figure 1, c). We incorporate the unlabeled data through two SSL formulations. In the neural framework, consistency regularization encourages two independently perturbed views of the same enzyme-substrate representation, whether labeled or unlabeled, to produce similar predictions (Figure 1, d). In the graph framework, labeled and unlabeled observations are represented as nodes in a nearest-neighbor graph, and Laplacian regularization encouraged strongly connected nodes to have similar inferred kinetic values (Figure 1, e).

**Figure 1.**
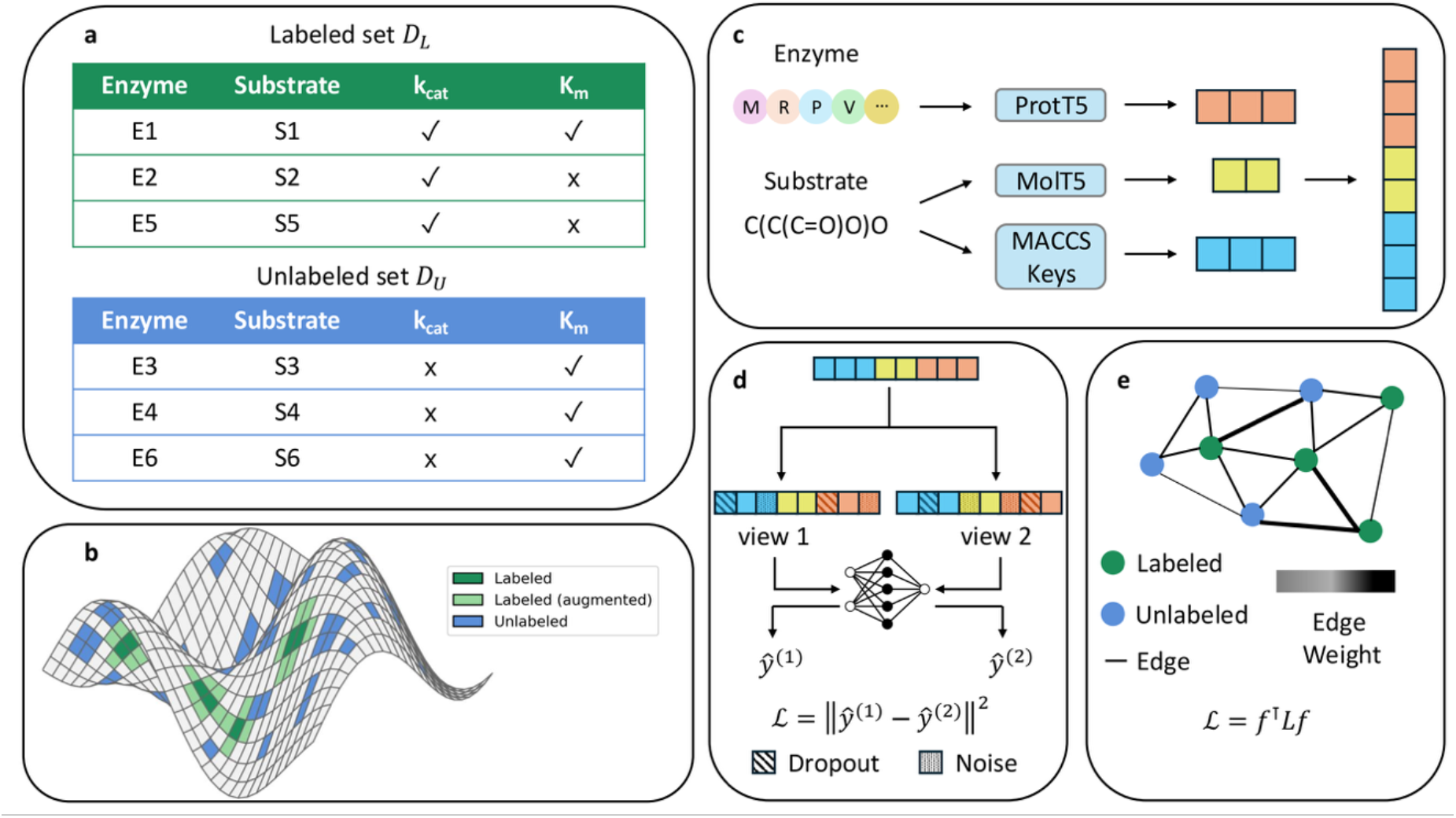
Overview of the semi-supervised learning framework for enzyme kinetic prediction. (a) Construction of labeled and unlabeled enzyme-substrate datasets. Pairs with experimentally measured *k*_cat_ values constitute the labeled set whereas those lacking *k*_cat_ but containing another reported kinetic parameter, such as *K*_m_, are treated as unlabeled observations. (b) Conceptual illustration of the labeled and unlabeled observations within the enzyme-substrate feature manifold. (c) Enzyme and substrate representations used for prediction. Enzyme sequences are encoded using ProtT5 embeddings while substrates are represented using MolT5 embeddings and MACCS fingerprints. The resulting features are concatenated into a joint enzyme-substrate representation. (d) Neural consistency regularization: two independently perturbed views of each labeled or unlabeled enzyme-substrate representation are generated using dropout and feature noise, and the model is encouraged to produce consistent predictions across the two views. (e) Graph-Laplacian regularization: labeled and unlabeled observations form nodes in a nearest-neighbor graph, with edge weights reflecting similarity in feature space. The Laplacian penalty encourages connected observations to have similar predicted kinetic values while supervised loss is applied only to labeled observations.

### 2.1 Unlabeled records expand enzyme coverage within the *k*cat prediction domain

The labeled CataPro dataset comprised 27,658 enzyme-substrate records spanning a broad range of log_10_(*k*_cat_) values with a median of 0.62 and an interquartile range of -0.4 to 1.54 (Figure 2, a). The unlabeled set was made up of the 16,187 enzyme-substrate pairs with a reported *K*_m_ but no corresponding *k*_cat_ in CataPro. To characterize the relationship between labeled and unlabeled data, we examine their feature space distribution. The unlabeled records overlapped with the labeled data in the first two principal components of the joint enzyme-substrate representation (Figure 2, b). However, the two principal components only explained 13.5% of the total variance. Hence, we quantified this overlap relative to the actual prediction setting by comparing, within each fold, the distance of the test set and unlabeled data to the same labeled training set (Figure 2, c). The unlabeled records were generally closer to the training data than the test set observations with median nearest-neighbor distances of 13.22 and 15.67, respectively. Moreover, 96.5 ± 1% of the unlabeled data fell within the distance envelope containing 95% of the test set. Therefore, the unlabeled pool largely occupies the same representation region in which model generalization is evaluated, rather than constituting a separate out-of-distribution dataset. This makes it a plausible source of additional information for SSL without requiring the model to extrapolate into a different feature space. The unlabeled dataset primarily expanded the enzyme rather than the substrate space: 71.9% of records paired a previously unseen enzyme sequence with a substrate already represented in the labeled dataset, while 15.3% contained both a previously observed enzyme and substrate (Figure 2, d). Overall, these results show that the unlabeled data extend enzyme sequence coverage while remaining closely connected to the supervised prediction domain, providing a natural source of information for *k*_cat_ prediction.

**Figure 2.**
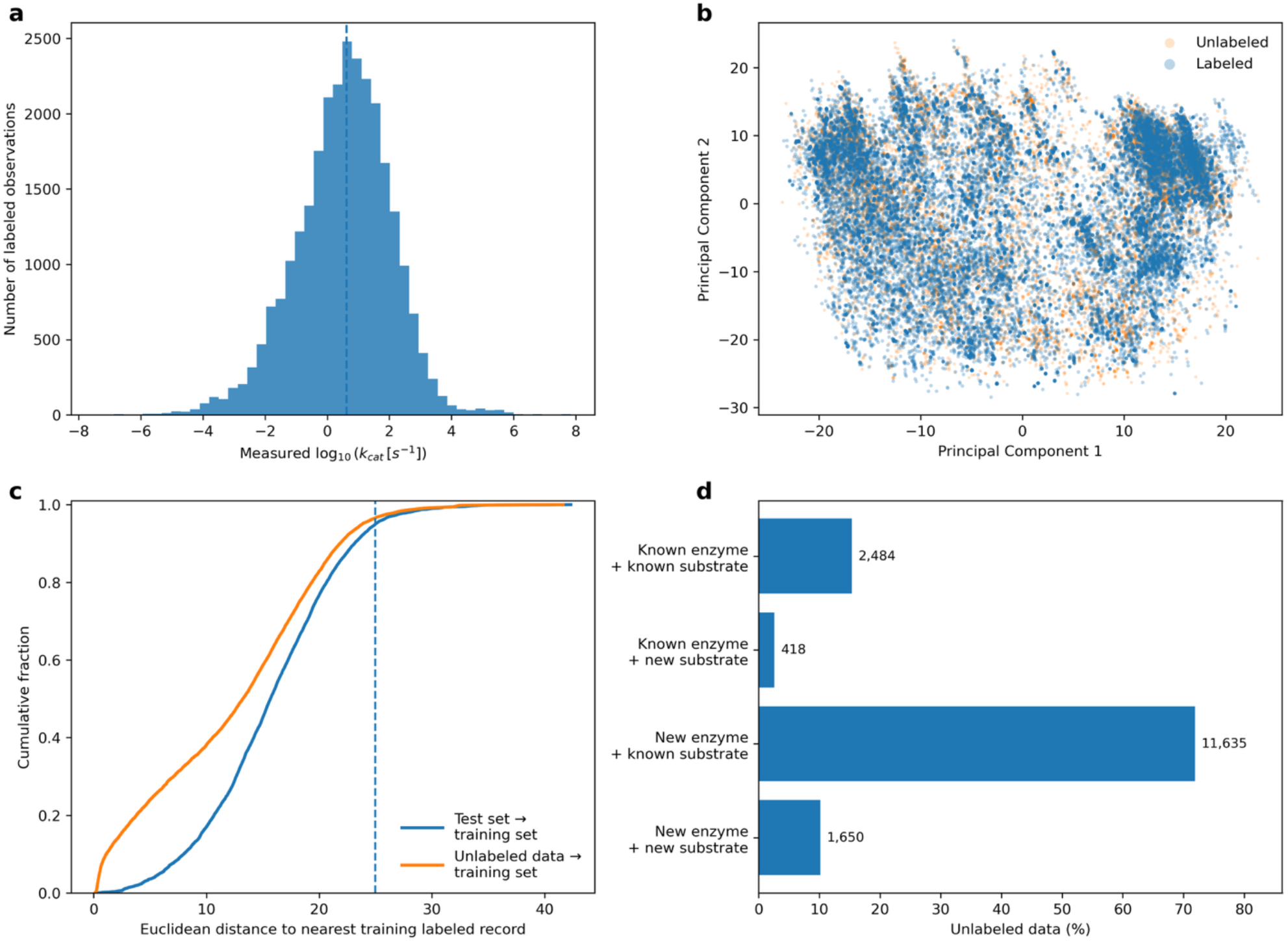
Distribution and characteristics of labeled and unlabeled records. (a) Distribution of measured log_10_(*k*_cat_) values across the labeled enzyme-substrate dataset from CataPro. (b) Principal component analysis (PCA) of the standardized joint enzyme-substrate representations for labeled and unlabeled records; Principal Component 1 and Principal Component 2 explain 8.2% and 5.3% of the total variance respectively. (c) Cumulative distributions of the Euclidean distance from test and unlabeled records to their nearest labeled training observation, calculated within each fold. The dashed line indicates the pooled 95^th^ percentile of labeled test-to-training distances. (d) Composition of the unlabeled dataset according to whether the enzyme sequence and substrate were previously observed in the labeled dataset. Bar labels indicate the number of records in each category.

### 2.2. Unlabeled records and consistency regularization improve *k*cat predictions

We first evaluated whether consistency regularization improved *k*_cat_ prediction relative to conventional supervised training and, critically, whether the addition of unlabeled enzyme-substrate data provided a benefit beyond consistency regularization applied to labeled observations alone. The three training configurations were evaluated using the ten pre-defined CataPro folds clustered at 40% sequence identity, with five independent random seeds per fold, yielding 50 train/test runs for each configuration. For statistical comparisons, predictions were averaged across seeds within each outer fold, resulting in ten paired observations per training condition.

Consistency regularization on labeled observations substantially improved the supervised baseline, reducing mean RMSE from 1.425 to 1.352 and increasing mean PCC from 0.447 to 0.476 (Figure3, a, b). It is worth noting that the baseline performance was comparable to what was reported in CataPro^7^, with differences arising due to differences in model implementation. Incorporating the 16,187 unlabeled enzyme-substrate records produced additional improvement, further reducing RMSE to 1.339 and increasing mean PCC to 0.479 relative to labeled consistency alone (Figure 3, a, b and Table S1). Importantly, the additional improvement from unlabeled data was consistent across the sequence-clustered test folds (Figure 3). After averaging across random seeds within each fold, the combined model (i.e., the model regularized using both labeled and unlabeled sequences) outperformed labeled consistency in all 10 folds for RMSE and *R*^2^, with significant paired differences by Wilcoxon signed-rank testing (*p* ≈ 0.002 for both metrics). However, PCC and Spearman’s correlation showed little systematic difference between the two consistency-based models. The parity plots similarly showed that all three models retained compression toward the center of the *k*_cat_ distribution, with overprediction at low values and underprediction at high values. However, consistency regularization produced a more concentrated high-density core with reduced dispersion of predictions, particularly in the combined model, while reducing several large prediction errors for moderate *k*_cat_ observations (Figure 3, e-g). These patterns suggest that the improvement in predictive accuracy was associated primarily with reduced prediction variability and extreme errors, rather than recovery of the full dynamic range of *k*_cat_ values.

**Figure 3.**
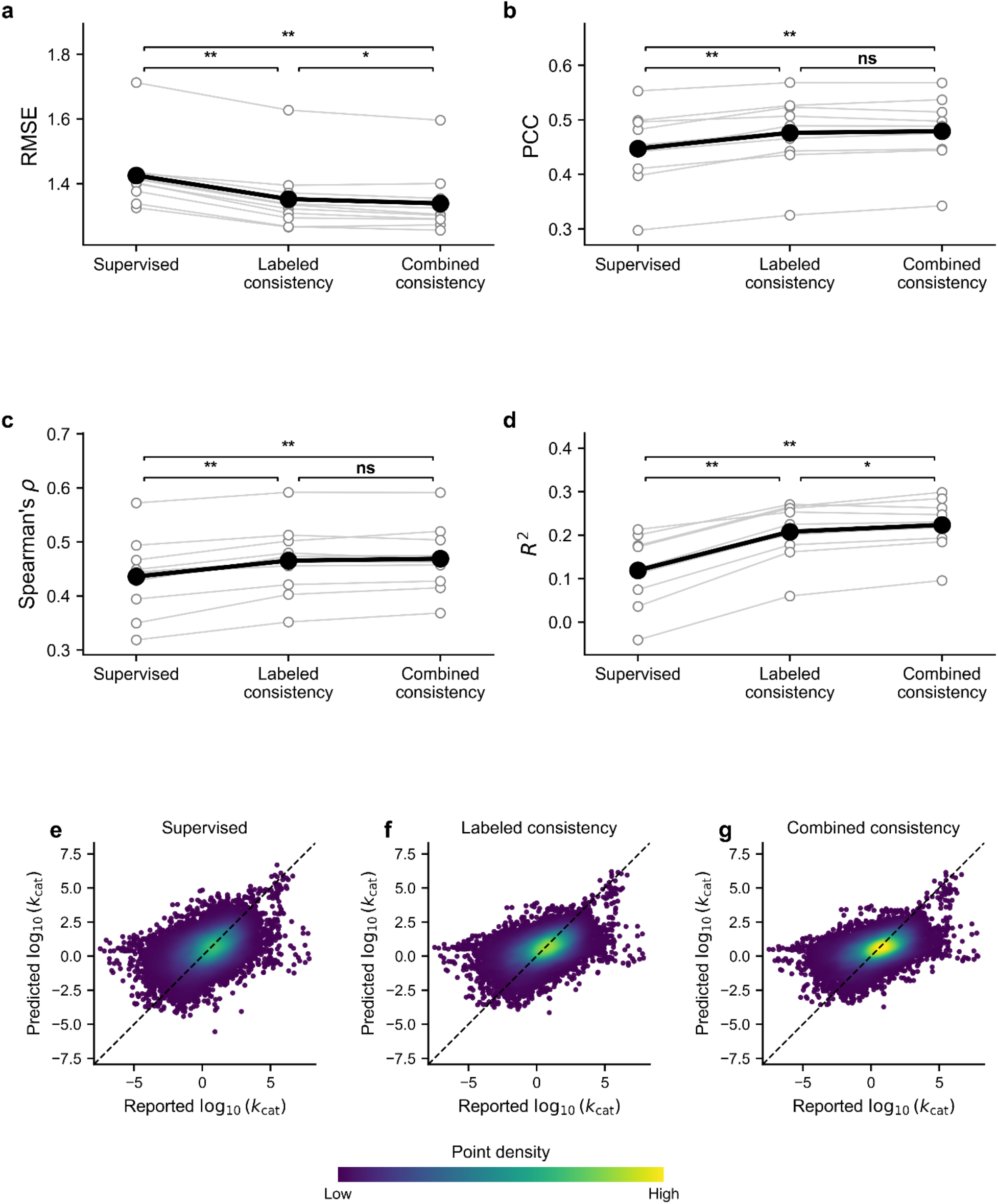
Performance of the supervised baseline, labeled-consistency model, and combined consistency model across the ten sequence-clustered folds from CataPro. Panels show (a) root-mean-square error (RMSE), (b) Pearson correlation coefficient (PCC), (c) Spearman rank correlation coefficient *ρ*, and (d) coefficient of determination *R*^2^. For each training condition, open gray circles represent the performance of an individual outer fold after averaging across five independent random seeds. Thin gray lines connect results from the same outer fold across training conditions. Large black circles and the connecting black line indicate the mean performance across the ten outer folds. Brackets denote pairwise comparisons between the different training conditions. \**p* < 0.05, \*\**p* < 0.01, and ns = not significant. Panels (e-g) show reported versus predicted log_10_(*k*_cat_) values for the (e) supervised, (f) labeled-consistency, and (g) combined-consistency models, respectively. Predictions for each enzyme-substrate observation were averaged across the five random seeds before visualization. Dashed lines indicate the line of identity.

The effect of unlabeled consistency was therefore most evident in magnitude-sensitive metrics of the ML task, including quadratic error and explained variance, rather than in the correlation or rank ordering of enzyme-substrate pairs. This indicates that the unlabeled records primarily improved the numerical estimates of *k*_cat_, rather than substantially rearranging which observations were predicted to have higher or lower turnover rates.

### 2.3. SSL benefits only emerge as labeled supervision becomes sufficient

We next examined whether the contribution of unlabeled enzyme-substrate records depends on the amount of labeled *k*_cat_ data available during training. The labeled training set was progressively reduced from 100% to 1% of its original size, while the full unlabeled pool remained available to the combined model (Figure 4).

**Figure 4.**
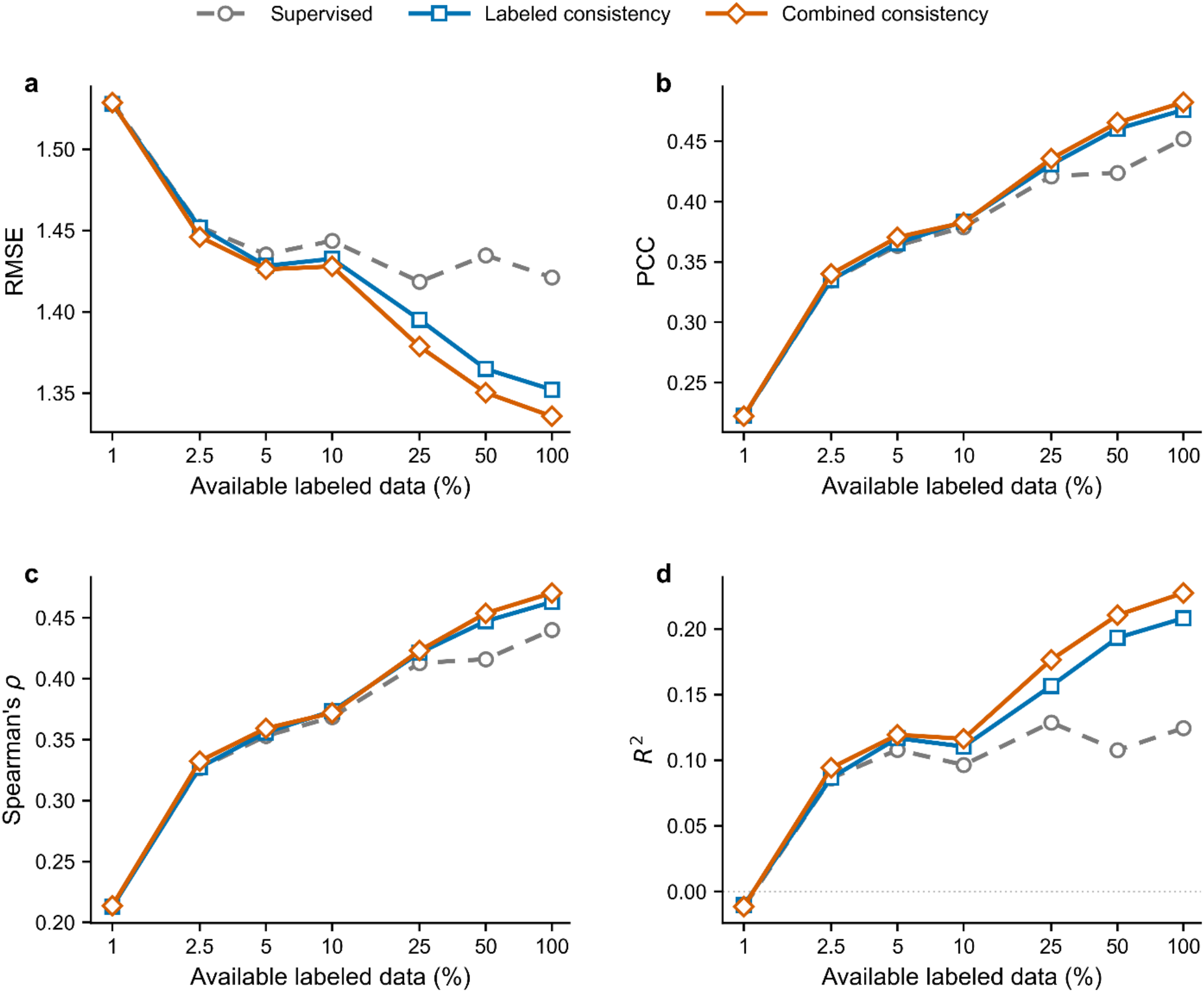
Effect of the availability of labeled data on *k*_cat_ prediction under supervised and consistency-regularized training. Curves show (a) root-mean-square error (RMSE), (b) Pearson’s correlation coefficient (PCC), (c) Spearman’s rank coefficient (*ρ*) and (d) coefficient of determination (*R*^2^) for the supervised baseline, labeled-consistency model, and combined consistency model. The fractions of available labeled data were calculated from the available training pool while 16,187 unlabeled records were used for each fraction.

At the most extreme scarcity condition, corresponding to only 1% of the labeled data, all three training strategies performed similarly and yielded a weak predictive signal. As additional labeled data became available, consistency-based training increasingly separated from the supervised baseline. The combined model generally performed best, with its advantage over labeled consistency becoming most apparent at moderate-to-high labeled fractions. For example, at 25% labeled-data availability, RMSE decreased from 1.419 for supervised training to 1.395 with labeled consistency and 1.377 with combined consistency, with this ordering persisting at higher labeled fractions (Figure 4, a).

The PCC, Spearman’s *ρ*, and *R*^2^ trends mirrored those observed for RMSE (Figure 4, b-d). These results indicate that unlabeled records do not substitute for the supervised signal required to learn the kinetic target. Instead, their benefit emerges once sufficient labeled data are available to establish the underlying *k*_cat_ relationship, after which consistency regularization on both labeled and unlabeled enzyme-substrate pairs provides further improvement in predictive accuracy.

### 2.4 Performance saturates with increasing the pool size of unlabeled data

We tested whether the improvement from SSL increased with the size of the unlabeled dataset. The effect of unlabeled consistency was evident across 25%, 50%, and 100% labeled-data regimes,but the magnitude of improvement did not scale proportionally with the size of the unlabeled pool (Figure 5). At 50% and 100% labeled-data availability, substantial gains were obtained after introducing up to 5,000 unlabeled enzyme-substrate records, after which performance largely plateaued or showed modest non-monotonic variation. For example, with all labeled data available, RMSE decreased from 1.352 with no unlabeled data to 1.336 with 5,000 unlabeled records (Figure 5, a), while PCC increased from 0.476 to 0.482 (Figure 5, b). In contrast, under the 25% labeled-data regime, performance continued to improve gradually as the unlabeled pool increased, with RMSE decreasing from 1.395 to 1.376 across the full unlabeled-data range. Similar patterns were observed for PCC, Spearman’s *ρ*, and *R*^2^ (Figure 5, c, d). These results indicate that the amount of unlabeled data required to approach maximal benefit depends on the amount of labeled supervision available. Relatively small unlabeled pools were sufficient when labeled data were abundant whereas gains accumulated more gradually under greater label scarcity.

**Figure 5.**
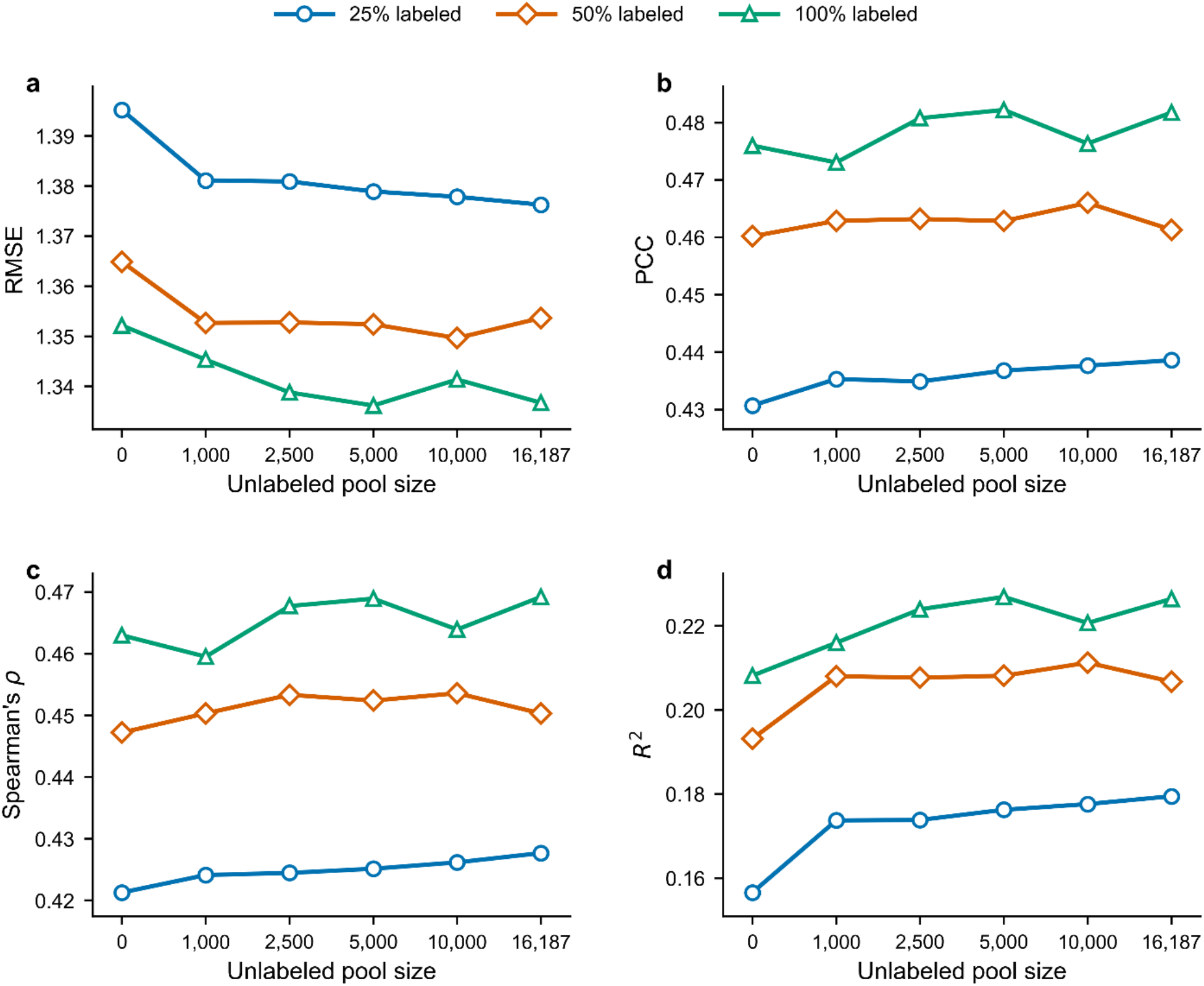
Predictive performance as a function of unlabeled pool size assessed using (a) root-mean-square error (RMSE), (b) Pearson’s correlation coefficient (PCC), (c) Spearman’s rank coefficient (*ρ*) and (d) coefficient of determination (*R*^2^). These models were run using 27,658 labeled data points. For 0 unlabeled pool size, the model is equivalent to labeled-only consistency regularization.

### 2.5. Unlabeled data also improves graph-based regression

To determine whether the benefit of unlabeled enzyme-substrate records was specific to neural consistency regularization, we evaluated a graph-based semi-supervised regression framework using the same CataPro representation and sequence-clustered test folds. *k*-Nearest-neighbor (*k*NN) was used as the supervised baseline because its neighborhood-based prediction provides a natural counterpart to the local graph structure used for Laplacian regularization. After hyperparameter optimization (Table S2), graph regularization using labeled data alone improved performance relative to supervised *k*NN, reducing RMSE from 1.509 to 1.482 and increasing Spearman’s *ρ* from 0.373 to 0.381 (Figure 6, a, c and Table S1). Incorporating the 16,187 unlabeled enzyme-substrate records into the graph produced a substantially larger improvement, reaching a mean RMSE of 1.414 and increasing Spearman’s *ρ* to 0.411 (Figure 6, a, c and Table S1). The combined Laplacian-*k*NN model outperformed labeled Laplacian-*k*NN in all ten folds for RMSE and *R*^2^, with each comparison significant by paired Wilcoxon signed-rank test ( *p* = 0.00195). Improvements were also observed in nine of ten folds for PCC and Spearman *ρ* (*p* = 0.00586 for both) (Figure 6). The parity plots showed a progressive concentration of predictions within a more compact high-density core from supervised kNN to labeled and combined Laplacian regularization, accompanied by fewer extreme prediction errors (Figure 6, e-g). Thus, the advantage of incorporating unlabeled records was reproduced using a non-neural graph-based SSL mechanism, indicating that the observed benefit is not specific to perturbation-based consistency training.

**Figure 6.**
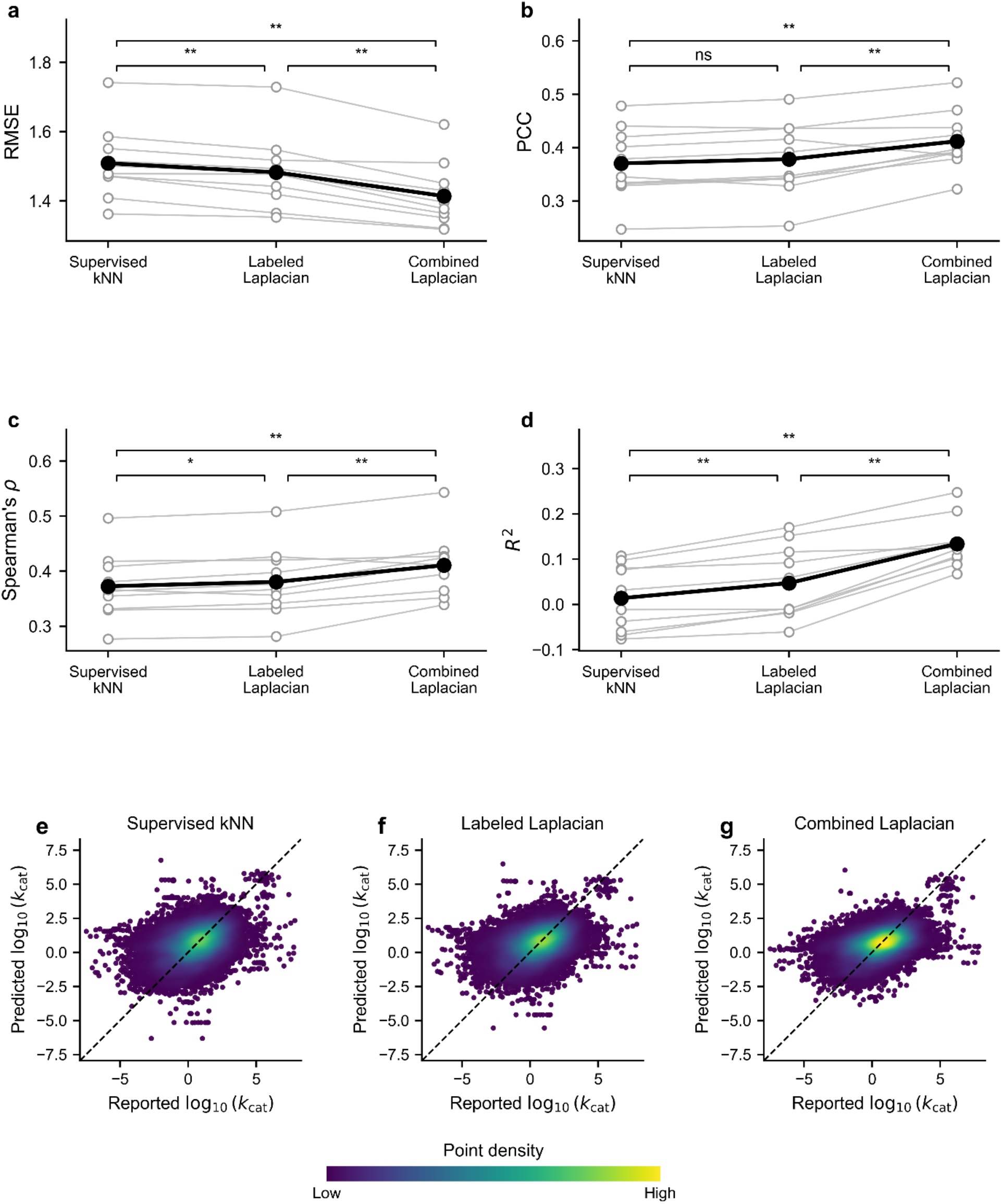
Performance of the supervised *k*NN baseline, labeled Laplacian-graph model, and combined Laplacian-graph model across the ten sequence-clustered folds from CataPro. Panels show (a) root-mean-square error (RMSE), (b) Pearson correlation coefficient (PCC), (c) Spearman’s rank correlation coefficient *ρ*, and (d) coefficient of determination (*R*^2^). For each training condition, open gray circles represent the performance of an individual outer fold after averaging across five independent random seeds. Thin gray lines connect results from the same outer fold across training conditions. Large black circles and the connecting black line indicate the mean performance across the ten outer folds. Brackets denote pairwise comparisons between the different training conditions. \**p* < 0.05, \*\**p* < 0.01, and ns = not significant. Panels (e-g) show reported versus predicted log_10_(*k*_cat_) values for the (e) supervised, (f) labeled-Laplacian, and (g) combined-Laplacian models, respectively. Predictions for each enzyme-substrate observation were averaged across the five random seeds before visualization. Dashed lines indicate the line of identity.

The combined graphs showed substantial integration between labeled and unlabeled observations, with labeled-unlabeled edges accounting for approximately 32% of graph edges on average across seed-fold runs (Table S3). Only 0.57% of unlabeled observations, on average, belonged to connected components containing no labeled node, indicating that nearly all unlabeled observations were connected directly or indirectly to experimentally labeled regions of the graph (Table S3). These diagnostics support the ability of the Laplacian objective to propagate information between labeled and unlabeled enzyme-substrate observations.

Hyperparameter selection also differed between the labeled and combined graph formulations (Table S2). The labeled Laplacian model more frequently selected broader graph neighborhoods whereas the combined model favored smaller neighborhoods and more often selected the lower Laplacian regularization weight. This shift is consistent with the addition of unlabeled observations altering local graph connectivity and reducing the need for more aggressive smoothing across larger neighborhoods. Prediction-neighborhood size remained concentrated at small values in both settings, indicating that the main effect of the unlabeled data was on graph construction and signal propagation rather than on the final local interpolation step.

### 2.6. Unlabeled data utility varies across feature space

We investigated whether the contribution of unlabeled enzyme-substrate records depended on their location within the feature space. Unlabeled data were first grouped according to their proximity to the labeled training data within each fold, based on the feature representations. Specifically, they were ranked by their mean Euclidean distance to the 20 nearest labeled training observations and divided into four quartiles, from Q1 (closest) to Q4 (farthest). Each quartile contained approximately 4,047 unlabeled observations. A randomly sampled quartile of the unlabeled pool was included as a size-matched reference, along with the complete set of 16,187 unlabeled observations.

For neural consistency regularization, performance varied only modestly across proximity quartiles (Figure 7, a). Mean RMSE values were ∼1.34 for all quartiles which was also consistent with using the whole unlabeled set. Hence, neural SSL performance did not exhibit a monotonic dependence on the proximity of unlabeled observations to the labeled training data. Graph-based SSL showed greater sensitivity to the proximity to training data (Figure 7, b). Mean RMSE decreased from 1.487 for Q1 to 1.466 for Q2 and 1.449 for Q3 before increasing to 1.464 for Q4. The random quartile subset reached an RMSE of 1.438 while the complete unlabeled pool produced the best graph-based performance at 1.414. Similar trends were observed for PCC (Figure S1, a, b), Spearman’s *ρ* (Figure S2, a, b), and *R*^2^ (Figure S3, a, b). These results indicate that proximity to labeled observations alone did not identify the most informative unlabeled subset.

**Figure 7.**
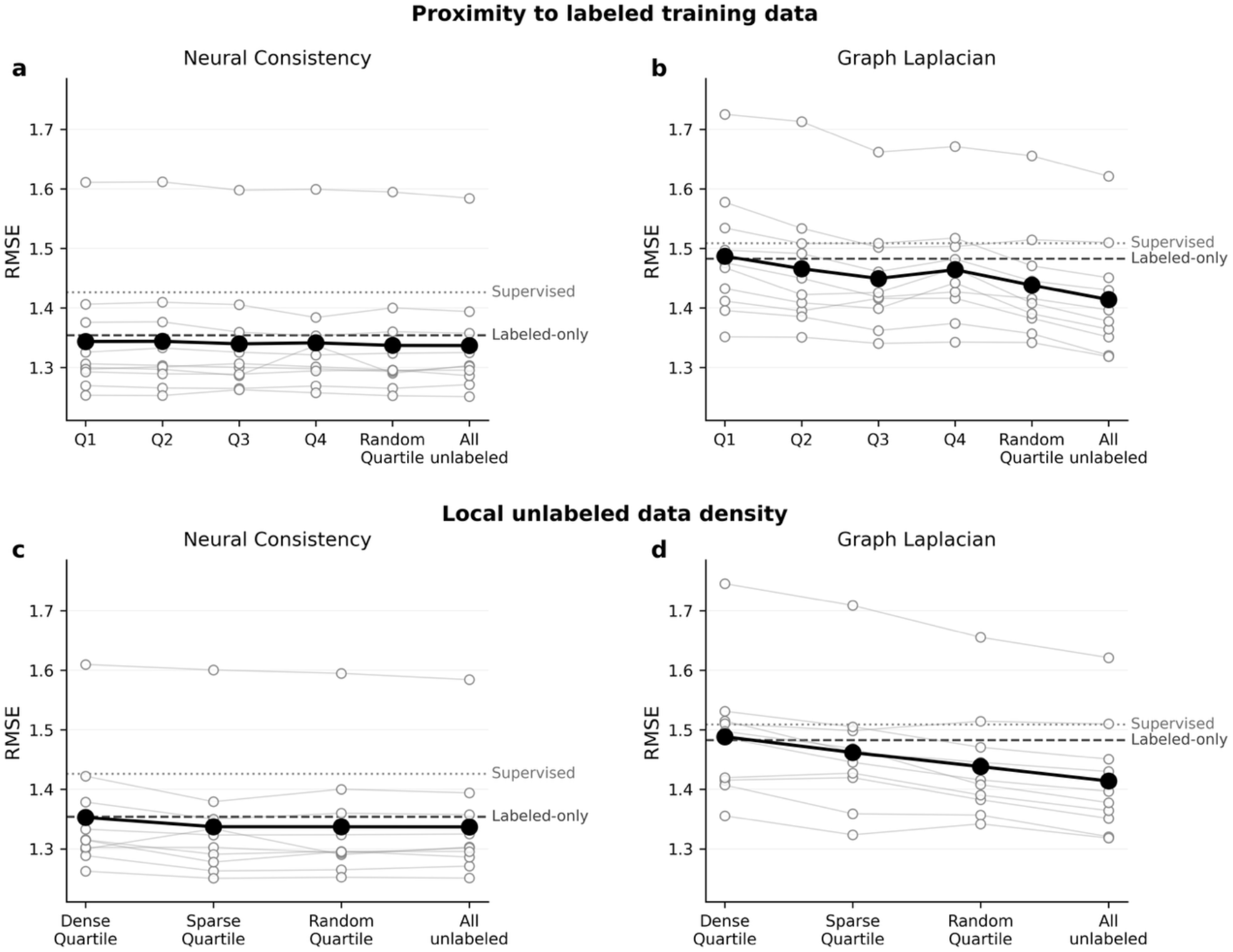
Dependence of SSL performance on the feature representation space of unlabeled data using RMSE as a metric. (a, b) Unlabeled observations were ranked by their mean Euclidean distance to the 20 nearest labeled training observations and divided into four approximately equal proximity quartiles, from Q1 (closest) to Q4 (farthest). Neural consistency is shown in (a) and graph Laplacian regularization in (b). (c, d) Local unlabeled data density was quantified using the mean Euclidean distance to the 20 nearest unlabeled observations. The dense quartile contained the 25% of records with the smallest mean neighbor distances whereas the sparse quartile contained the 25% with the largest distances. Neural and graph results are shown in (c) and (d), respectively. A randomly sampled quartile and the complete set of 16,187 unlabeled observations were included as reference conditions. Open gray circles represent individual fold performances after averaging across five random seeds, with thin gray lines connecting results from the same fold. Large black circles and connecting lines indicate the mean across the ten folds. Dashed horizontal lines indicate the labeled consistency or regularization condition while dotted lines indicate the corresponding supervised baseline.

Thereafter, we examined whether local density within the unlabeled dataset was associated with model performance. For each unlabeled observation, local density was quantified by the mean Euclidean distance to its 20 nearest unlabeled neighbors. The 25% of observations with the smallest mean neighbor distances were designated the dense subset whereas the 25% with the largest distances formed the sparse subset.

For the neural model, the dense subset did not provide any improvement over labeled consistency (Figure 7, c). In contrast, the sparse subset decreased mean RMSE from 1.354 to 1.337. Sparse observations outperformed dense observations in 8 of 10 outer folds for RMSE (*p* = 0.048; Figure 7, c), PCC (*p* = 0.027; Figure S1, c) and *R*^2^ (*p* = 0.037; Figure S3, c). Also, it surpassed denser observation in 7/10 outer folds for Spearman’s *ρ* (*p* = 0.027; Figure S2, c). Performance with the sparse subset was comparable to both the random quartile and the complete unlabeled pool. A stronger density dependence was observed for graph-based SSL (Figure 7, d). The dense subset yielded a mean RMSE of 1.488, which fell short compared to the labeled Laplacian regularization, yet the sparse subset improved RMSE to 1.462. Sparse observations outperformed dense observations in 8 of 10 outer folds for RMSE and *R*^2^, with *p* = 0.0098 for each comparison (Figure 7, d and Figure S3, d). The random quartile subset further decreased RMSE to 1.438, and the complete unlabeled pool reached 1.414. The random subset outperformed the sparse subset in 8 of 10 folds for *R*^2^ (*p* = 0.027), while the complete pool outperformed the random subset in all 10 folds (*p* = 0.002). The correlation metrics followed a similar trend (Figure S1, d and Figure S2, d). These results demonstrate that the predictive contribution of unlabeled observations varied with their representation-space distribution, with densely populated regions contributing less benefit than sparse or more broadly sampled subsets.

### 2.7. SSL gains extend across wild-type and mutant enzymes

We examined whether the improvements associated with SSL were restricted to wild-type enzymes or were also retained for mutants. The labeled dataset contained 16,828 wild-type and 10,830 mutant records while the unlabeled dataset had 11,994 wild-type and 4,193 mutant entries. Prediction performance was therefore evaluated separately for wild-type and mutant enzyme-substrate records within the CataPro test folds.

For the neural model, consistency regularization improved prediction in both subgroups relative to conventional supervised training (Table 1). Among wild-type observations, RMSE improved from 1.385 to 1.299 and 1.279 and *R*^2^increased from 0.098 to 0.206 and 0.231, respectively. The same overall trend was observed for mutant enzymes, although absolute prediction performance was lower. Mutant RMSE decreased from 1.486 to 1.432 with labeled consistency and to 1.429 with the combined model while *R*^2^ increased from 0.065 to 0.133 and 0.137, respectively. The additional effect of the unlabeled data was therefore more pronounced for wild-type observations than for mutants. For mutant observations, PCC and Spearman correlation were essentially unchanged following the addition of unlabeled data and were marginally higher under labeled consistency. This mirrors the overall CataPro results, in which the contribution of unlabeled consistency was expressed more strongly through magnitude-sensitive error metrics than through changes in rank ordering.

**Table 1.** Performance of the neural consistency and graph Laplacian regularization across wild-type (WT) and mutant CataPro observations under supervised, labeled, and combined regularization conditions. Bold values indicate the best performing condition within each subgroup for each metric. Statistical significance was assessed using two-sided paired Wilcoxon signed-rank tests across the 10 splits. Asterisks indicate significant improvement relative to the supervised baseline (*p < 0.05, **p < 0.01, ***p < 0.001) whereas daggers indicate significant improvement relative to labeled-only conditions (†p < 0.05, ††p < 0.01, †††p < 0.001). Significance symbols are shown in parentheses beneath the corresponding values.

| Framework | Labeled Test Set Subgroup | Condition | RMSE | $R^2$ | PCC | $\rho$ |
| --- | --- | --- | --- | --- | --- | --- |
| Neural Network | WT | Supervised | 1.385 ± 0.079 | 0.098 ± 0.099 | 0.449 ± 0.085 | 0.432 ± 0.084 |
| | | Labeled consistency | 1.299 ± 0.076<br>(**) | 0.206 ± 0.080<br>(**) | 0.486 ± 0.080<br>(**) | 0.464 ± 0.081<br>(**) |
|  |  | Combined consistency | **1.279 ± 0.074**<br>(**,††) | **0.231 ± 0.076**<br>(**,††) | **0.496 ± 0.076**<br>(**,††) | **0.473 ± 0.079**<br>(**,††) |
| | Mutant | Supervised | 1.486 ± 0.163 | 0.065 ± 0.092 | 0.424 ± 0.076 | 0.419 ± 0.079 |
| | | Labeled consistency | 1.432 ± 0.161<br>(**) | 0.133 ± 0.083<br>(**) | **0.443 ± 0.080**<br>(**) | **0.444 ± 0.075**<br>(**) |
|  |  | Combined consistency | **1.429 ± 0.149**<br>(**) | **0.137 ± 0.071**<br>(**) | 0.440 ± 0.071 | 0.442 ± 0.071 |
| *k*-Nearest Neighbor | WT | Supervised | 1.416 ± 0.070 | 0.057 ± 0.078 | 0.411 ± 0.068 | 0.390 ± 0.074 |
|  |  | Labeled Laplacian | 1.394 ± 0.076<br>(**) | 0.086 ± 0.090<br>(**) | 0.417 ± 0.072 | 0.398 ± 0.077<br>(*) |
|  |  | Combined Laplacian | **1.343 ± 0.079**<br>(**,††) | **0.153 ± 0.070**<br>(**,††) | **0.438 ± 0.066**<br>(*) | **0.419 ± 0.073**<br>(*,†) |
|  | Mutant | Supervised | 1.643 ± 0.158 | -0.144 ± 0.118 | 0.304 ± 0.089 | 0.330 ± 0.073 |
|  |  | Labeled Laplacian | 1.613 ± 0.155<br>(**) | -0.102 ± 0.105<br>(**) | 0.313 ± 0.079 | 0.340 ± 0.066 |
|  |  | Combined Laplacian | **1.521 ± 0.126**<br>(**,††) | **0.020 ± 0.059**<br>(**,††) | **0.358 ± 0.060**<br>(*,†) | **0.381 ± 0.058**<br>(*,††) |

The graph-based framework showed a stronger additional effect of unlabeled data in both subgroups (Table 1). For wild-type observations, RMSE decreased from 1.416 with supervised *k*NN to 1.394 with labeled Laplacian-*k*NN and 1.343 with combined Laplacian-*k*NN, accompanied by improvements in *R*^2^, PCC, and Spearman’s *ρ*. The effect was particularly pronounced for mutant observations. Mean PCC improved from 0.304 under supervised conditions to 0.358 under the combined model while RMSE decreased from 1.643 to 1.521. Thus, the benefit of incorporating unlabeled observations was not restricted to wild-type enzymes and, in the graph framework, was particularly evident for mutant predictions.

### 2.8. SSL generalizes to a low data adenylate kinase landscape

We next evaluated the same SSL strategies on an independent adenylate kinase (ADK) dataset to investigate whether the benefit of unlabeled sequence information extends to local dense datasets with a few labeled points. In contrast to CataPro, this dataset represents an extremely low data setting, comprising 175 experimentally characterized ADKs with adenosine diphosphate (ADP) as a substrate. We used the 30 previously defined train/test partitions in Muir et al. (2025), with 140 labeled enzymes used for training and 35 held out for evaluation in each split. Moreover, the authors had retrieved an additional 4,974 diverse ADK sequences from bacteria and archaea lacking kinetic measurements, which we utilized as our unlabeled pool^2^. Because the size of the 1,280-dimensional ESM-2 representations^24^ was large relative to the labeled training set, the embeddings were reduced using PCA fitted exclusively to the labeled training data within each split. We retained 32 principal components for the neural model and 50 components for the graph-based model. Models were evaluated independently for *k*_cat_, *K*_m_, and *k*_cat_/*K*_m_. For single target prediction using ESM-2, our supervised baseline results were consistent with those reported in the original paper using different ML models, with Spearman’s *ρ* ranging from 0.38 to 0.46 for *k*_cat_ and 0.13 to 0.23 for *K*_m_^2^ (Tables S4 and S5).

The most pronounced SSL benefit was observed for *k*_cat_/*K*_m_ (Table 2). SSL produced a modest improvement in the neural consistency model, but larger numerical gains in the graph-based model, particularly in where it increased *R*^2^ from 0.009 to 0.135. Notably, incorporation of the unlabeled ADKs also reduced the variability in *R*^2^ across the 30 splits, suggesting more robust performance across different selections of labeled training enzymes. The overall result was consistent across the remaining kinetic parameters: improvements for *k*_cat_ were modest (Table S4) while *K*_m_ benefited primarily from the graph-based formulation (Table S5).

**Table 2.**
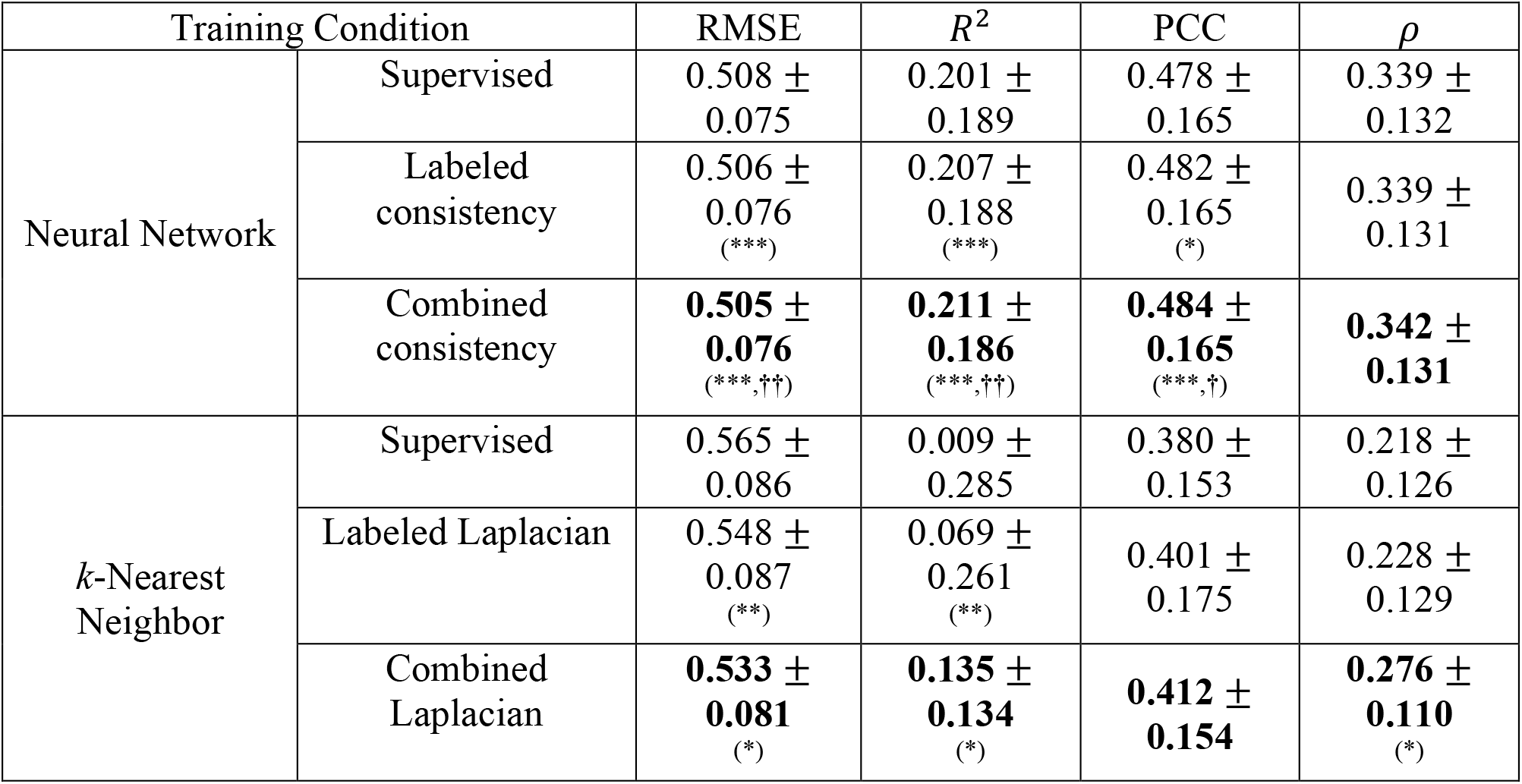
Predictive performance for *k*_cat_/*K*_m_ in the adenylate kinase dataset. Performance is reported for the neural consistency and graph-Laplacian frameworks under supervised, labeled regularization, and combined labeled–unlabeled conditions across the 30 predefined ADK train/test splits. Values are mean ± standard deviation across splits. Bold values indicate the best performing condition within for each metric. Statistical significance was assessed using the paired Wilcoxon signed-rank test across the 30 splits. Asterisks indicate significant improvement relative to the supervised baseline (*p < 0.05, **p < 0.01, ***p < 0.001) whereas daggers indicate significant improvement relative to the corresponding labeled-only regularization condition (†p < 0.05, ††p < 0.01, †††p < 0.001). Significance symbols are shown in parentheses beneath the corresponding values.

## 3. Discussion

The central finding of this study is that leveraging unlabeled enzyme-substrate data alongside labeled data during task-specific training through regularization-based SSL approaches can improve kinetic parameter prediction without generating any pseudo-labels. Across the CataPro *k*_cat_ benchmark, the addition of unlabeled records improved prediction using two mechanistically distinct SSL strategies: consistency regularization in a neural model and Laplacian regularization in a nearest-neighbor graph. The effect was reproduced across sequence-identity-clustered folds. These results suggest that experimentally documented enzyme-substrate observations for which the prediction target is unavailable can still contain useful information about the structure of the feature space. Importantly, however, the label scarcity analysis showed that this information does not replace labeled kinetic targets. At the lowest labeled data fractions, all models had little predictive ability and unlabeled observations provided minimal additional benefit. The contribution of SSL emerged only after the labeled data were sufficient to establish a meaningful relationship between the biochemical representation and the kinetic target. The SSL strategy was also applied to an independent ADK dataset representing a local low-data sequence landscape.

Previous work has shown that directly incorporating unlabeled protein sequences into regression is not uniformly beneficial. Olivares-Gil et al. evaluated semi-supervised wrapper methods, Co-Training and Tri-Training regressors, across 19 protein fitness datasets using homologous sequences as unlabeled observations and tested labeled training sizes ranging from the complete dataset down to 50 sequences^21^. The wrapper methods showed a Spearman’s *ρ* comparable to the supervised baseline, but none remained competitive as label availability decreased. These results suggest that assigning predicted targets to unlabeled homologs and iteratively incorporating them into the training set can be sensitive to the quality of the initial supervised model, particularly under severe label scarcity. In contrast, the present study incorporates unlabeled observations without assigning pseudo-labels. Instead, we utilize them to regularize either local prediction consistency or the geometry of a graph whose target values remain anchored exclusively by labeled kinetic data. The reproducible improvements achieved by our SSL algorithms across the ten predefined CataPro folds demonstrate consistent gains that prior pseudo-label-based wrapper methods have not achieved, opening new opportunities for SSL-driven protein engineering.

Our systematic analysis of neural regularization approaches helps separate two sources of improvement that can otherwise be conflated in SSL. Applying consistency regularization to labeled observations alone produced a substantial improvement over conventional supervised training, indicating that enforcing invariance to light perturbations of the enzyme and substrate representations is itself an effective regularizer. The subsequent addition of unlabeled observations produced a smaller but reproducible further improvement. This distinction is conceptually related to O^2^DENet, which recently showed that perturbation-based invariance can improve out-of-distribution enzyme kinetic prediction by generating altered versions of already labeled enzyme-substrate pairs. The present results support the utility of this local smoothness principle while showing that the effect is not exhausted by augmentation of labeled data: independent unlabeled enzyme-substrate observations provide additional support points at which the prediction function can be constrained. Our combined approach therefore obtains complementary information from augmentation and unlabeled data. These perturbations define which local changes the model should ignore around observations already represented in the labeled set whereas unlabeled observations determine additional locations in the feature space at which that smoothness constraint can be imposed.

The parity plots provide additional insight into how the improvement in performance is achieved. Across the three training conditions, predictions remain substantially compressed toward the center of the observed *k*_cat_ distribution, with the models tending to overpredict low *k*_cat_ values and underpredict high *k*_cat_ values. This compression becomes more apparent under consistency regularization, even as RMSE, *R*^2^, and, to a lesser extent, PCC improve. This behavior is consistent with the tendency of regression models trained on noisy and sparsely sampled targets to shrink predictions toward the mean when the available features do not provide sufficient information to confidently resolve extreme values^25,26^. In the present setting, the effect may be particularly pronounced because *k*_cat_ spans several orders of magnitude and the labeled data provide limited coverage of the extremes. Regularization encourages locally similar enzyme-substrate representations to produce similar predictions which can suppress highly variable predictions in sparsely supported regions of the feature space. The resulting reduction in extreme errors can improve RMSE and *R*^2^ even when the overall prediction range becomes narrower. Because PCC and Spearman’s *ρ* depend primarily on whether predictions preserve the relative ordering of observations rather than whether their magnitudes are perfectly calibrated, these metrics can also remain stable or improve despite this compression. Thus, the parity plots suggest that the principal benefit of SSL is not the recovery of the full dynamic range of *k*_cat_, but rather a reduction in large prediction errors.

The saturation of the neural SSL effect provides additional insight into this mechanism. Much of the improvement was recovered with around 5,000 unlabeled observations, and a randomly sampled quartile produced essentially the same final performance as the complete 16,187 record pool. Consistency training does not propagate numerical target estimates from one observation to another. Instead, each unlabeled example acts as a location at which two stochastic views are required to yield similar predictions. Because the neural network uses a single globally shared parameterization, gradients generated in one portion of the feature space modify the same function used elsewhere. Once a sufficiently diverse collection of data samples the relevant input distribution, additional examples become increasingly redundant. This also helps explain why the four proximity quartiles produced nearly indistinguishable neural performance, as distance from the labeled set is not itself the quantity optimized by consistency regularization. Rather, what matters more directly is whether the selected observations expose the network to regions in which its response to realistic perturbations can be usefully constrained.

The graph-based model behaved differently because unlabeled data is incorporated into the model structurally rather than simply through an additional loss evaluation. Each new unlabeled record becomes a graph node that alters local neighborhoods and edge weights by creating new paths linking different regions of the feature space. Hence, this produced a substantially larger additional effect of unlabeled data relative to labeled Laplacian regularization, despite the graph model generally having lower absolute predictive performance than the neural model, at least on the dataset used in this study. Nearly all unlabeled nodes were connected directly or indirectly to a component containing labeled data. Consequently, the graph continued to benefit from increasing the size and breadth of the unlabeled pool even after neural consistency had largely saturated. This contrast is useful because it shows that the same unlabeled data pool can contribute through different mechanisms: the neural model primarily uses them to sample locations at which to regularize a global function as opposed to graph regularization that uses them to modify the topology over which kinetic information is smoothed. We did not perform a separate labeled and unlabeled pool size sensitivity analysis for the graph framework. The neural label-scarcity experiments had already established that a minimum amount of labeled information is required before unlabeled data can provide a useful signal. For the graph model, nearly all unlabeled nodes were connected directly or indirectly to labeled components, with only approximately 0.5% remaining unanchored. We therefore used the full unlabeled pool, as subsampling was expected to reduce graph connectivity rather than reveal a large subset of uninformative observations.

The unlabeled data utility analysis further indicates that coverage, rather than simple closeness to labeled examples, dictates the observed benefit. Selecting unlabeled data closest to the labeled training data did not consistently improve either model, and the graph performed better with intermediate proximity groups than with the closest quartile. Similarly, records drawn from the densest regions of the unlabeled space contributed almost no improvement whereas the sparsest quartile provided a substantially larger benefit in both frameworks. A plausible explanation is redundancy: an unlabeled observation surrounded by many nearly equivalent labeled or unlabeled observations provides relatively little additional information about the biochemical feature distribution. By contrast, observations occupying less densely sampled spaces extend the regions over which smoothness or graph structure can be constrained. That being said, the performance of the random quartile and complete pool precludes a stronger conclusion that the sparsest observations are universally optimal. For neural consistency, sparse, random, and complete pools performed similarly, consistent with rapid saturation once broad coverage had been obtained. For the graph, the random subset outperformed the sparse subset and the complete pool performed best, suggesting that graph construction benefits from a combination of broad coverage and sufficient local sampling to maintain connectivity. Thus, selecting only isolated observations can extend the graph into new regions but may simultaneously remove intermediate nodes needed to represent continuous relationships between them. Nevertheless, these geometric analyses should be interpreted as associations rather than a causal decomposition of the value of unlabeled data. Density and proximity in the feature space are correlated with biochemical composition, and the dense and sparse subsets differed in their distributions of wild-type and mutant enzymes, enzyme classification (EC) annotations, and enzyme-substrate novelty. In addition, Euclidean neighborhoods in a 1,959-dimensional concatenated representation are only an approximation to biochemical similarity. The observed patterns therefore do not establish that low density itself causes improved SSL performance. Rather, they show that repeatedly sampling highly populated regions is insufficient and that useful unlabeled information is associated with broader feature space coverage.

The wild-type and mutant analyses further suggest that the effect of SSL is not restricted to a single type of enzyme sequence. Both subgroups benefited from regularization relative to supervised training, although the additional unlabeled contribution differed by model. In the neural model, the further improvement from unlabeled data was more evident for wild-type observations whereas the graph formulation produced a pronounced improvement for mutants, including a shift from negative to positive mean *R*^2^. One possible factor contributing to this difference may be the composition of the unlabeled pool, which contains substantially more wild-type than mutant observations and may therefore provide a more representative consistency signal for wild-type regions. Graph regularization can additionally benefit mutants through paths involving nearby wild-type and mutant nodes rather than requiring the unlabeled observations themselves to match the test set.

The independent ADK analysis supports the broader applicability of the framework while also illustrating its limitations. Muir et al. (2025) showed that the ADK catalytic landscape is highly rugged and that protein language model representations organize sequences more readily by structural features than by catalytic activity; incorporating experimental labels through ProteinNPT improved prediction relative to lightweight supervised models. Our results extend this broader picture by showing that unlabeled sequence information can also contribute to kinetic prediction when incorporated directly through task-specific consistency and graph-based regularization, rather than through ProteinNPT’s distinct prediction framework. Neural consistency produced only modest additional gains in this extreme low-data setting, but graph regularization produced larger improvements, particularly for *k*_cat_/*K*_m_. This is consistent with the CataPro label-scarcity results: when the labeled target information is extremely limited, smoothness constraints alone cannot reconstruct a rugged catalytic landscape. Meanwhile, the graph results suggest that even under such conditions, explicitly representing the geometry of thousands of unlabeled homologous sequences can stabilize prediction across the labeled landscape. That being said, due to the lack of biologically grounded enzyme-substrate pairs for ADK, the unlabeled pool consisted of homologous sequences without any corresponding experimental validation. Some sequences may provide less reliable information for the SSL task and may partly contribute to the smaller and more variable SSL gains observed in the low-data setting.

Overall, the results support a view of unlabeled data as more than unused entries awaiting kinetic measurement, demonstrating that they can provide valuable information when leveraged alongside labeled data through SSL. Once sufficient labeled data define the kinetic landscape, unlabeled observations can constrain how that relationship varies across the enzyme-substrate space. The extent of the resulting improvement depends on both the learning mechanism and the distribution of the unlabeled observations: neural consistency rapidly reaches saturation once the relevant space is adequately sampled while graph regularization continues to benefit from the increased coverage and connectivity of larger unlabeled pools. This distinction provides a practical basis for deciding not only whether unlabeled data should be incorporated into enzyme kinetic prediction, but also how their geometry and redundancy may determine their worth.

## 4. Methods

### 4.1. Problem formulation

We formulated the enzyme kinetic prediction task as a semi-supervised regression over experimentally characterized enzyme-substrate pairs. Let an enzyme-substrate observation be denoted by,

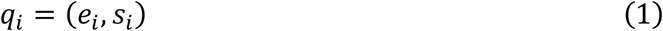

where *e*_*i*_ is an enzyme amino acid sequence and *s*_*i*_ is the corresponding substrate.

For prediction of a kinetic target, the available data were separated into a labeled set *D*_*L*_ and an unlabeled set *D*_*U*_,

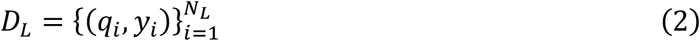

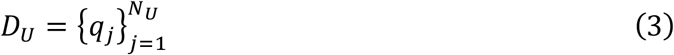

where *y*_*i*_ is the experimentally measured kinetic parameter for each observation in *D*_*L*_ . The objective is to learn a regression function *f*_*θ*_: *χ* → ℝ parametrized by *θ*, using both *D*_*L*_ and *D*_*U*_, while never assigning pseudo-labels to the latter.

### 4.2. Kinetic dataset

The present study used the *k*_cat_ dataset and sequence-clustered cross-validation folds introduced with CataPro. The original CataPro datasets were assembled from BRENDA release 2023.1 and SABIO-RK v2.20. During CataPro curation, entries with ambiguous units were removed and repeated annotations corresponding to the same enzyme sequence-substrate pair were collapsed by retaining the maximum reported *k*_cat_ and minimum reported *K*_m_. The resulting data contained 27,658 unique *k*_cat_ entries, 42,018 unique *K*_m_ entries, and 25,831 enzyme-substrate pairs for which both parameters were reported. CataPro further clustered enzyme sequences at a sequence-similarity threshold of 0.4 and assigned the resulting clusters among ten partitions for cross-validation^7^.

In this study, as a labeled set, we focused on the *k*_cat_ dataset from CataPro for the prediction task with a regression target of,

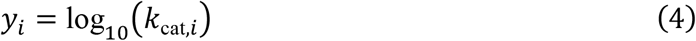

The unlabeled set was obtained from the experimentally characterized enzyme-substrate pairs with a reported *K*_m_ but no corresponding *k*_cat_, yielding 16,187 entries.

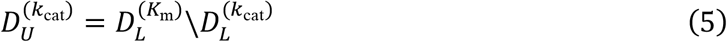

The existence of a *K*_m_ measurement was used only to identify the corresponding enzyme-substrate combination as a valid biochemical reaction. Thus, for each *q*_*j*_ ∈ *D*_*U*_, the model had access to the enzyme and substrate representations but not to any estimate of *k*_cat,j_.

#### 4.3. Feature representation and standardization

Each enzyme-substrate pair was represented using the feature definition employed by CataPro. The enzyme sequence *e*_*i*_ was encoded using a 1024-dimensional ProtT5-XL-UniRef50 embedding, **p**_*i*_ ∈ ℝ^1024^, while substrate *s*_*i*_ was represented using both a 768-dimensional MolT5 embedding, **m**_*i*_ ∈ ℝ^768^, and a 167-dimensional MACCS molecular fingerprint, **c**_*i*_ ∈ ℝ^167^. The three representations were concatenated to form a feature vector **x**_*i*_, as shown in Equation 6.

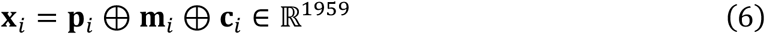

Feature standardization was performed separately within every outer fold and random seed experiment to avoid information transfer from the validation or test sets. For a feature coordinate *d*, the mean and standard deviation were estimated using only the labeled training set as,

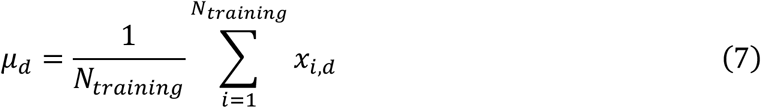

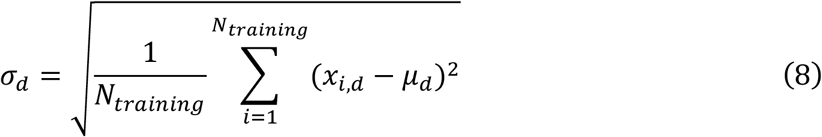

Each feature was subsequently transformed as,

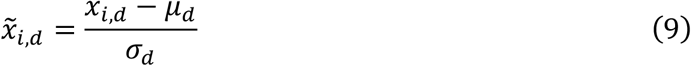

The standardized feature vector was denoted as 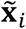.

### 4.4. Neural consistency-regularized regression

#### 4.4.1. Regression model

To isolate the effect of the semi-supervised learning objective, a fixed shallow multilayer perceptron (MLP) was used for all experiments. The network consisted of **x**_*i*_ as a 1959-dimensional input layer, one hidden layer containing 256 nodes with rectified linear unit activation, dropout with a probability of 0.2, and a single linear regression output.

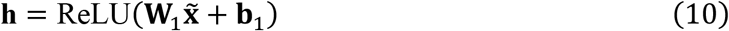

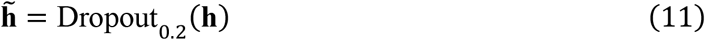

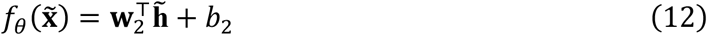

where *W*_1_ ∈ ℝ^256×1959^ and *b*_1_ ∈ ℝ^256^ . The parameter set *θ* = {**W**_1_, **b**_1_, **w**_2_, *b*_2_} was optimized jointly during training.

#### 4.4.2. Stochastic perturbation of enzyme-substrate representations

Consistency regularization requires multiple stochastic views of the same underlying observation. We therefore defined an augmentation operator *A*(**x**; **ξ**) parametrized by independently sampled random variables *ξ*. Perturbations were modality-specific because ProT5 and MolT5 components are continuous latent representations whereas MACCS keys are derived from a binary molecular fingerprint. The augmentation parameters were fixed throughout all experiments: Gaussian noise standard deviation of 0.02, 5% feature dropout for ProtT5 and MolT5, and 2% feature dropout for MACCS.

For a standardized input, 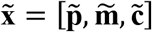, the augmented ProtT5 component would be,

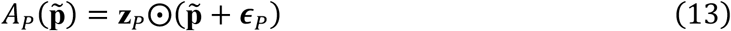

where *ϵ*_*p*_∼*N* (**0**, 0.02^2^ 734 **I**_1024_) and the individual elements of the dropout mask followed 735 z_*p*_∼Bernoulli(0.95).

The MolT5 representation was perturbed analogously,

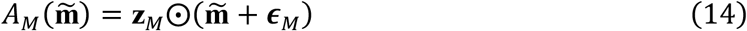

with *ϵ*_*p*_∼ *N* (**0**, 0.02^2^**I**_123_) and **z**_*p*_∼Bernoulli(0.95).

For the standardized MACCS coordinates, only a light feature-dropout was used.

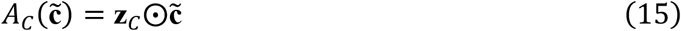

where **z**_*C*_ ∼Bernoulli(0.98).

Therefore, the complete augmented representation was,

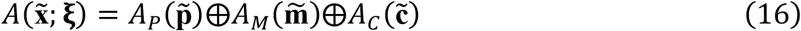

For every consistency calculation, two perturbations were sampled independently where 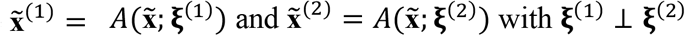.

#### 4.4.3. Supervised and consistency-regularized objectives

For a labeled batch 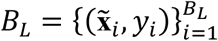,the supervised component was the mean squared error between the log-transformed *k*_cat_ values and predictions obtained from the unperturbed standardized representations,

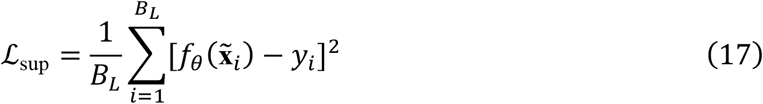

To distinguish the general regularizing effect of representation perturbation from the contribution of external unlabeled records, consistency regularization was first applied to labeled observations themselves. Two independent views of each labeled example were constructed, and the corresponding prediction discrepancy was minimized,

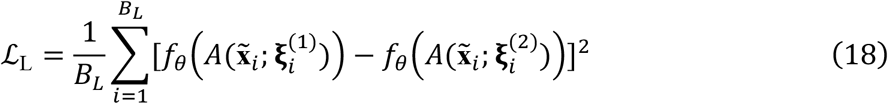

This term imposes local smoothness of the regression function around labeled enzyme-substrate observations.

For an unlabeled batch, 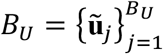, two independently perturbed views of each unlabeled enzyme-substrate representation were generated, and the unlabeled consistency loss was,

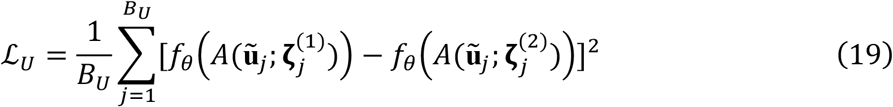

The complete learning objective was defined as,

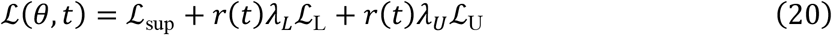

where *λ*_*L*_ and *λ*_*U*_ control labeled and unlabeled consistency respectively.

To prevent consistency objectives from dominating optimization before the regression function had acquired a supervised signal, their contribution was increased linearly over the first 30 epochs using a ramp function,

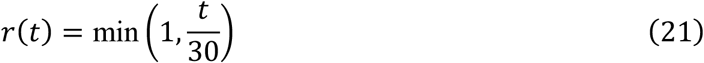

The principal comparison consisted of three objectives as presented in Table S6. The comparison between the supervised and labeled-consistency models quantifies the effect of perturbation-based regularization alone. The comparison between labeled consistency and combined consistency isolates the additional effect of introducing the unlabeled enzyme-substrate set.

#### 4.4.4. Sequence-clustered cross-validation and model training

Model performance was evaluated using the 10 predefined CataPro folds generated from enzyme-sequence clustering at a 0.4 similarity threshold. Let *F* = {*F*_1_, …, *F*_10_} denote these partitions. For outer fold *k*, let held-out test set be,

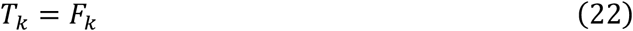

and the available training set be,

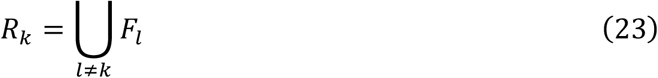

No observation from *T*_*k*_ was used for parameter estimation, early stopping, or model selection. For each random seed σ, *R*_*k*_ was further partitioned into 90% inner-training subset and a 10% inner-validation subset such that,

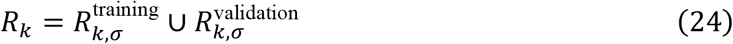

The inner-validation set was used only for early stopping. The standardization parameters in Equations 7 and 8 were estimated using 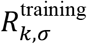 alone. The corresponding transform was subsequently applied to 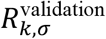, *T*_*k*_, and **u**. The experiment was repeated for five random seeds σ ∈ {42, 43, 44, 45, 46} across all ten outer folds, producing 50 train/test runs for each training condition.

Within each seed-fold combination, model initialization and labeled batch ordering were matched across the supervised, labeled-consistency, and combined-consistency. Models were optimized using Adam with learning rate of 10^−4^ and a batch size of 128. Training proceeded for at most 200 epochs. After each epoch, predictions were obtained for the unaugmented inner-validation data, and the validation root-mean-square error was calculated. Training was terminated if no improvement occurred for 20 consecutive epochs. Network dropout was active only during training and was disabled for validation and test predictions. These hyperparameters were held fixed for all three objectives.

### 4.5. Graph-based semi-supervised regression

To determine whether unlabeled enzyme-substrate records could improve prediction through a mechanism distinct from neural consistency regularization, we implemented a graph-based semi-supervised regression framework. The same labeled datasets, feature representations, and sequence-clustered outer folds used for the neural experiments were retained. Here, enzyme-substrate observations were represented as nodes in a nearest-neighbor graph, and their relationships were incorporated through graph-Laplacian regularization. Three configurations were evaluated to distinguish the effect of graph regularization from the additional contribution of unlabeled observations: (i) supervised *k*-nearest-neighbor regression using labeled observations only as a baseline, (ii) Laplacian-regularized *k*-nearest-neighbor regression using a graph containing labeled observations only, and (iii) combined Laplacian-regularized *k*-nearest-neighbor regression using a graph containing both labeled and unlabeled observations.

#### 4.5.1 Graph construction

For each training split, every enzyme-substrate observation was represented by its standardized feature vector 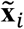 . Euclidean distance was used to define local neighborhoods. For a graph containing *N* observations, each node *i* was connected to its *k*_*g*_ nearest neighboring nodes. To accommodate variations in local feature-space density, edge weights were assigned using a locally scaled radial basis function. The local bandwidth σ_*i*_ was defined as the Euclidean distance between node *i* and its *k*_*g*_-th nearest neighbor. The directed weight between nodes *i* and *j* was then defined as,

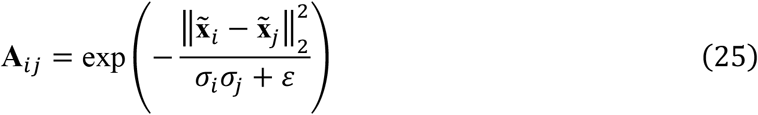

where *ε* is a small constant used to prevent division by zero. The weighted adjacency matrix **A** was symmetrized by retaining the larger weight for each connected node pair and self-connections were removed. The weighted degree matrix was defined by,

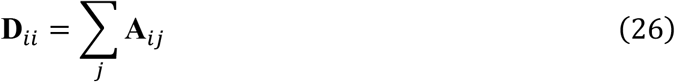

and the symmetric normalized graph Laplacian was calculated as,

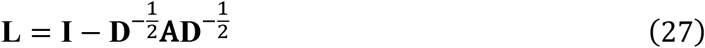

Graph construction was performed independently within each training fold. Held-out test sets were not included as graph nodes.

#### 4.5.2. Laplacian-regularized graph signal estimation

Let **f** = (*f*_1_, …, *f*_*N*_) ^⊤^ denote the regression signal defined over the graph nodes. For labeled nodes, the corresponding *k*_cat_ labels were available whereas the unlabeled nodes contained no *k*_cat_ target. Graph signal estimation balanced fidelity to the target values with smoothness over the graph such as,

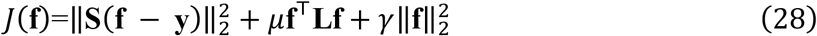

Here, *μ* controls graph smoothness, *γ* is a small stabilization term, and **S** is a diagonal label-indicator matrix defined as,

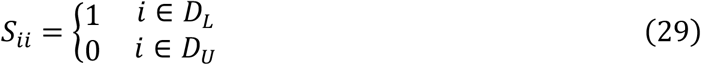

The graph smoothness term **f**^⊤^**Lf** penalizes large differences in the inferred regression signal between strongly connected observations. The resulting objective function has a linear solution as shown in Equation 30.

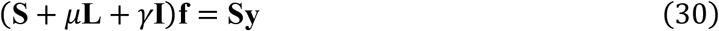

The system was solved using a sparse conjugate-gradient solver, with a least-squares sparse solver used as a fallback if convergence was not achieved. The stabilization coefficient *γ* was fixed at 10^−4^.

#### 4.5.3. Inductive prediction for the held-out test set

Because held-out test sets were excluded from graph construction, prediction was performed inductively after graph-signal estimation. For a test observation with feature vector 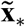, its *k*_*p*_ nearest graph nodes were identified in the same standardized feature space. The predicted target was calculated using inverse-distance weighting as,

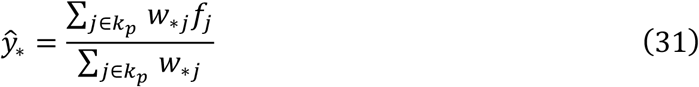

where

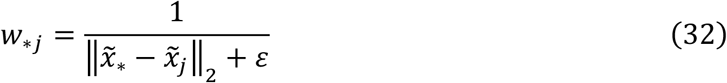

and *f*_*j*_ is the estimated graph signal associated with node *j*.

For supervised *k*-nearest-neighbor regression, the same prediction rule was applied using only labeled training observations, with *f*_*j*_ = *y*_*j*_.

#### 4.5.4. Graph model configurations

Similar to the neural regression setup, three graph-related training configurations were evaluated:

1. Supervised *k*NN: The model contained labeled observations only and used their target log_10_(*k*_cat_) values directly for inverse-distance-weighted *k*-nearest-neighbor prediction. No graph-smoothing objective was applied.
2. Labeled Laplacian-*k*NN: A graph was constructed using only labeled training observations. The graph signal was estimated using the Laplacian-regularized objective described above and the held-out test set was subsequently predicted from the smoothed labeled graph signal.
3. Combined Laplacian-*k*NN: The graph contained both labeled and unlabeled enzyme-substrate records. Target fidelity was imposed only on labeled nodes while both labeled and unlabeled nodes contributed to graph smoothness. Prediction was subsequently performed using the graph signal estimated over the combined node set.

This design enabled the contribution of graph regularization itself to be distinguished from the additional contribution of incorporating unlabeled enzyme-substrate observations.

#### 4.5.5. Hyperparameter selection and final model refitting

Graph-model hyperparameters were selected independently within each training fold. The labeled training data were first divided into a 90% inner-training subset and a 10% inner-validation subset using the corresponding random seed. Feature standardization was fitted exclusively to the labeled inner-training subset and applied to the inner-validation set and unlabeled data. Hyperparameters were selected by minimizing inner-validation RMSE. The prediction neighborhood size was evaluated over *k*_g_ ∈ {1, 3, 5, 10, 20, 40}, the graph neighborhood size over *k*_*p*_ ∈ {10, 20, 40, 80}, and the Laplacian regularization coefficient over *μ* ∈ {0.01, 0.1, 1, 10} . The stabilization coefficient *γ* was fixed at 10^−4^. After selecting the optimal hyperparameters, the inner validation split was returned to the training data. The feature scaler was refitted using the complete labeled training fold, and the final labeled or combined graph was reconstructed using the selected hyperparameters. The resulting model was evaluated on the corresponding held-out test fold. The graph experiments used the same ten sequence-clustered outer folds as the neural experiments and were repeated using random seeds 42, 43, 44, 45, and 46. Performance was therefore evaluated over 50 seed-fold runs before seed averaging for statistical analysis.

### 4.6. Label-scarcity analysis

To examine if the usefulness of unlabeled data depends on the amount of labeled data, a label-scarcity analysis was conducted using seed 42. The same ten outer test folds were retained so that decreasing the size of the labeled set affected only the data available for model training and optimization. For each outer fold *k*, the 90/10 inner-training/validation partition was first generated exactly as in the full-data experiment. Reduced labeled subsets were then constructed at *α* = 1%, 2.5%, 5%, 10%, 25%, 50% and 100% fractions. Importantly, the subsampling operation was applied independently to both the fixed inner-training and fixed inner-validation pools. This avoided retaining an artificially large validation set when only a small fraction of labels was assumed available.

Let Ω denote the indices of either the full inner-training or inner-validation pool and let *π*_Ω_ denote a fixed random permutation of its elements. For labeled fraction *α*, the number of retained observations was,

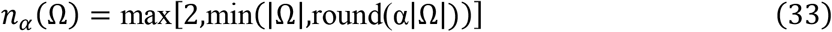

The reduced set was then defined from the first *n*_*α*_ positions of the fixed ordering,

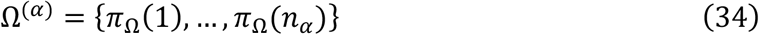

Consequently, for *α*_1_ < *α*_2_, the subset obeyed 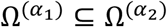, allowing differences across labeled fractions to be interpreted without confounding them with independently redrawn subsets. Separate fixed permutations were used for the training and validation pools. The full target-unlabeled pool *D*_*U*_ remained available at every labeled fraction. Feature scaling was re-estimated using only the reduced inner-training set at each value of *α*. The scarcity analysis was intended as a controlled sensitivity experiment and was therefore performed using a single fixed seed rather than included in the five seed significance analysis.

### 4.7. Unlabeled pool size analysis

A second sensitivity analysis evaluated the dependence of model performance on the number of unlabeled data. This experiment was performed using seed 42 at representative labeled fractions *α* of 25%, 50%, and 100%. The exact nested labeled subsets generated for the label-scarcity analysis were reused. The evaluated unlabeled pool sizes were 0, 1000, 2500, 5000, 10000, and 16187 (all *D*_*U*_). To ensure that differences between pool sizes reflected the addition of observations rather than independently sampled unlabeled sets, a single random permutation was generated as earlier to guarantee that Ω^(1000)^ ⊆ ⋯ ⊆ Ω^(16187)^.

### 4.8. Unlabeled data utility analysis

To understand the utility of unlabeled enzyme–substrate observations, we performed two complementary subset analyses based on the proximity of unlabeled data to labeled training observations and the local density within the unlabeled pool. All distances were calculated independently within each test fold using the same standardized 1,959-dimensional enzyme-substrate representation used for model training. For the proximity analysis, the Euclidean distance from each unlabeled observation to the labeled training set was calculated, and the mean distance to its 20 nearest labeled training observations 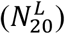 was used as its proximity score,

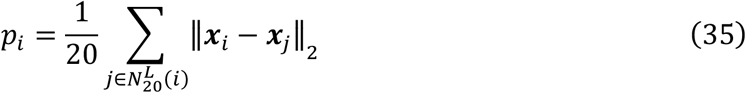

Unlabeled records were ranked according to *p*_*i*_ and divided into four approximately equal quartiles. Q1 contained the observations with the smallest proximity scores and was therefore closest to the labeled training data while Q4 contained those with the largest scores and was farthest from the labeled data. Ultimately, Q1–Q3 contained 4,047 observations each and Q4 contained 4,046 observations. For the density analysis, local density was defined relative to other unlabeled observations. For each unlabeled observation, the mean Euclidean distance to its 20 nearest unlabeled neighbors 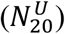 was calculated as,

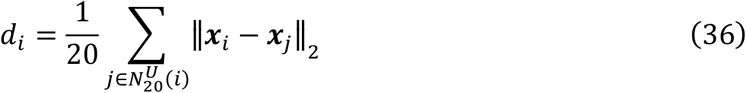

Smaller values of *d*_*i*_ corresponded to more densely populated regions of the unlabeled feature space. The 25% of observations with the smallest *d*_*i*_ values formed the dense subset (4047 records) whereas the 25% with the largest values formed the sparse subset (4046 records).

A randomly selected 25% subset of the complete unlabeled pool (4047 records) was included as a reference. Random subsets were generated independently for each random seed using fixed random states. The full unlabeled pool (16,187 records) and the corresponding labeled-only model were retained as additional reference conditions.

For neural consistency regularization, each unlabeled subset replaced the complete unlabeled pool while all network architecture, augmentation, optimization, and early stopping procedures were otherwise unchanged. For graph-based regression, the selected unlabeled subset was incorporated as graph nodes together with the labeled training observations. To isolate the effect of the unlabeled pool composition from differences in model selection, graph hyperparameters for the subset analyses were held fixed to those selected using the complete unlabeled pool for the corresponding seed and test fold. All subset experiments were repeated across the ten predefined CataPro test folds and five random seeds. Performance was first averaged across the five seeds within each test fold before statistical analysis.

### 4.9. Adenylate kinase dataset and evaluation protocol

We applied the same neural consistency and graph Laplacian frameworks to an independent previously reported adenylate kinase (ADK) dataset^2^. The dataset comprised 175 experimentally characterized ADK sequences with measurements of *k*_cat_, *K*_m_, and *k*_cat_/*K*_m_. We used the 30 predefined train/test partitions from the original study with 140 labeled ADKs used for model training and 35 held out for testing in each split. An additional set of 4,974 diverse bacterial and archaeal ADK sequences lacking kinetic measurements was used as the unlabeled pool. Each ADK sequence was represented using a 1,280-dimensional ESM-2 protein embedding^24^. Because the dimensionality of the feature vectors was large relative to the number of labeled training observations, principal component analysis (PCA) was applied separately within each split using only the labeled training data. The fitted transformation was then applied to the corresponding unlabeled and test sequences. 32 principal components were retained for the neural consistency model and 50 components for the graph-based model. The three kinetic targets *k*_cat_, *K*_m_, and *k*_cat_/ *K*_m_ were modeled independently. Performance was evaluated using the same regression metrics as for the CataPro experiments. Five random seeds (42-46) were used.

The neural regressor consisted of a single hidden layer with 32 units, ReLU activation, dropout of 0.1, and a linear output. Models were optimized with AdamW using a learning rate of 3 × 10^−4^, weight decay of 10^−3^, a labeled batch size of 32, and an unlabeled batch size of 128. Training proceeded for at most 300 epochs with early stopping after 30 epochs without sufficient validation improvement. Consistency regularization used two independently perturbed views of each representation. Gaussian noise with standard deviation 0.01 and feature dropout of 0.02 were applied to both labeled and unlabeled embeddings. The labeled-and unlabeled-consistency weights were both set to 0.25 and linearly ramped over the first 30 epochs. We used a smaller consistency weight given the much smaller labeled dataset. For each split, the best training epoch was selected according to validation RMSE.

For the graph-based regressor, the supervised baseline used inverse-distance-weighted *k*-nearest-neighbor regression on labeled ADKs only. The labeled-only condition constructed a graph from labeled ADKs and estimated a Laplacian-regularized regression signal. The combined condition constructed the graph from both labeled and unlabeled ADKs, while target fidelity was imposed only on labeled nodes. The graph hyperparameter sets were retained as is from the CataPro experiment and were selected by minimizing validation RMSE.

### 4.10. Evaluation metrics

Predictions were evaluated against experimental values of log_10_(*k*_cat_). For a test set containing *n* observations with target values *y*_*i*_ and predictions *ŷ*_*i*_, root-mean-square-error was calculated as,

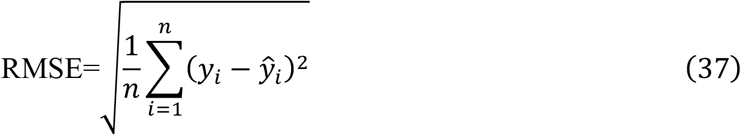

The coefficient of determination was,

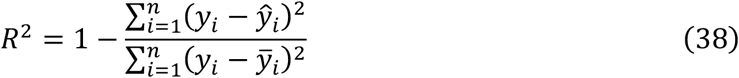

where

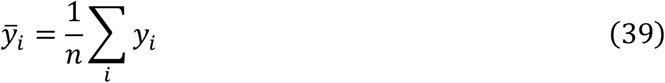

Linear association was quantified using Pearson’s correlation coefficient (PCC),

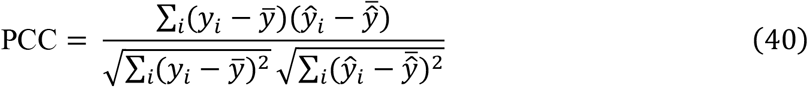

Rank agreement was evaluated using Spearman’s correlation coefficient,

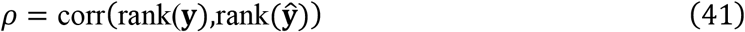

The same four metrics were computed independently for every seed-fold run.

### 4.11. Statistical analysis

Because the five random seeds share the same held-out fold, the 50 seed-fold runs were not treated as 50 independent observations for the primary statistical analysis. Instead, performance was first averaged over the five random seeds within each outer fold. For metric *v*, fold *k*, and training condition *c*, the average performance across was,

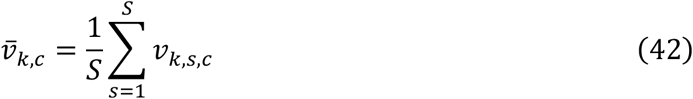

where *S* is the total number of random seeds used.

This produced ten paired observations per training condition (i.e., supervised, labeled, and combined), corresponding to the ten outer sequence-identity clustered folds. For a comparison between reference condition *a* and alternative condition *b*, paired fold-level differences were oriented such that positive values represented improvement. For *v* ∈ {RMSE},

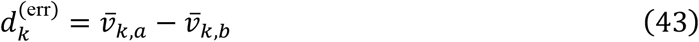

whereas for *v* ∈ {*R*^2^, PCC, *ρ*}, larger values were favorable,

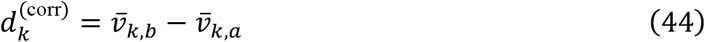

For the neural consistency experiments, comparisons included the pairwise combinations among supervised training, labeled consistency, and combined consistency. The same paired procedure was applied to the supervised *k*NN, labeled Laplacian-*k*NN, and combined Laplacian-*k*NN configurations. Because distributional assumptions are less secure with only ten outer folds, a two-sided Wilcoxon signed-rank test was utilized. The number of folds in which each alternative improved was additionally reported as a measure of directional consistency. A nominal significance threshold of *p* < 0.05 was used.

### 4.12. Implementation and reproducibility

Models were implemented in Python v3.11.5 using PyTorch. Data manipulation and numerical operations used pandas and NumPy. Preprocessing and regression metrics used scikit-learn. PCC,*ρ*, paired t, and Wilcoxon statistics were calculated using SciPy. Training was performed on an NVIDIA L40S GPU using New York University’s Torch High Performance Computing (HPC) cluster. Random states were explicitly controlled for Python, NumPy, and PyTorch as mentioned in Section 3.7. Exact outer fold assignments, inner splits, reduced labeled subsets, unlabeled subsets, per-run predictions, and per-run evaluation metrics were retained for reproducibility.

## Data and code availability

All data and code supporting the results of this study can be found in the article, supporting information, and the public repository available at https://github.com/ali-malli/SSL

## Acknowledgements

This work was supported by the New York University Research Catalyst Grant and the Korea Evaluation Institute of Industrial Technology [Project KEIT/MOTIE-20008416]. This work was also supported in part through the NYU IT High Performance Computing resources, services, and staff expertise.

## Author contributions

A. Malli conceptualized the study, designed the methodology, developed the algorithms, performed data analysis, conducted the evaluations, prepared visualization, and wrote the original draft of the manuscript. D.V. assisted with developing the algorithms and data analysis. A. Majumder assisted with data analysis and prepared visualization. J.R.K. supervised the project, secured the funding, and revised the original draft.

## Notes

The authors declare no competing interest.

