## Supplementary Information for "Going beyond labels: Semi-supervised learning for enzyme kinetic parameter prediction"

**Supplementary Information – List of Contents**

Table S1 – Performance of neural and graph-based supervised and semi-supervised models on the CataPro $k_{\mathrm{cat}}$ benchmark. Values are mean ± standard deviation across ten sequence-clustered folds after averaging across five random seeds within each fold. Bold values indicate the best performing condition for each metric.

Table S2 – Hyperparameter selection frequencies for graph-based regression. Each entry is the number of seed-fold runs selecting the value with the percentage of 50 runs in parentheses.

Table S3 – Connectivity properties of the final Laplacian graphs. Values are mean ± standard deviation across 50 final refitted graphs (10 folds over 5 random seeds).

Table S4 – Predictive performance for $k_{\mathrm{cat}}$ in the adenylate kinase dataset. Performance is reported for the neural consistency and graph-Laplacian frameworks under supervised, labeled regularization, and combined labeled–unlabeled conditions across the 30 predefined ADK train/test splits. Values are mean ± standard deviation across splits. Bold values indicate the best performing condition within for each metric.

Table S5 – Predictive performance for $K_{\mathrm{m}}$ in the adenylate kinase dataset. Performance is reported for the neural consistency and graph-Laplacian frameworks under supervised, labeled regularization, and combined labeled–unlabeled conditions across the 30 predefined ADK train/test splits. Values are mean ± standard deviation across splits. Bold values indicate the best performing condition within for each metric.

Table S6 – Final objective functions of the training conditions used in the neural consistency regularization experiment

Figure S1 – Dependence of SSL performance on the feature representation space of unlabeled data using PCC as a metric.

Figure S2 – Dependence of SSL performance on the feature representation space of unlabeled data using Spearman's $\rho$ as a metric.

Figure S3 – Dependence of SSL performance on the feature representation space of unlabeled data using $R^2$as a metric.

Table S1 – Performance of neural and graph-based supervised and semi-supervised models on the CataPro $k_{cat}$ benchmark. Values are mean ± standard deviation across ten sequence-clustered folds after averaging across five random seeds within each fold. Bold values indicate the best performing condition for each metric. Statistical significance was assessed using two-sided paired Wilcoxon signed-rank tests across the 10 splits. Asterisks indicate significant improvement relative to the supervised baseline (*p < 0.05, **p < 0.01, ***p < 0.001) whereas daggers indicate significant improvement relative to labeled-only regularization (†p < 0.05, ††p < 0.01, †††p < 0.001). Significance symbols are shown in parentheses beneath the corresponding values.

| Training condition | | RMSE | $R^2$ | PCC | $\rho$ |
|---|---|---|---|---|---|
| Neural Network | Supervised | 1.425 ± 0.107 | 0.119 ± 0.079 | 0.447 ± 0.070 | 0.436 ± 0.072 |
| | Labeled consistency | 1.352 ± 0.105 (**) | 0.208 ± 0.064 (**) | 0.476 ± 0.067 (**) | 0.465 ± 0.066 (**) |
| | Combined consistency | **1.339 ± 0.099** (**,†) | **0.224 ± 0.058** (**,†) | **0.479 ± 0.061** (**) | **0.469 ± 0.061** (**) |
| $k$-Nearest Neighbor | Supervised | 1.509 ± 0.104 | 0.014 ± 0.073 | 0.371 ± 0.067 | 0.373 ± 0.060 |
| | Labeled Laplacian | 1.482 ± 0.107 (**) | 0.047 ± 0.081 (**) | 0.378 ± 0.069 | 0.381 ± 0.062 (*) |
| | Combined Laplacian | **1.414 ± 0.094** (**,††) | **0.134 ± 0.054** (**,††) | **0.412 ± 0.055** (**,††) | **0.411 ± 0.057** (**,††) |

Table S2 – Hyperparameter selection frequencies for graph-based regression. Each entry is the number of seed-fold runs selecting the value with the percentage of 50 runs in parentheses.

| Parameter | Candidate values | Supervised $k$NN | Labeled Laplacian-$k$NN | Combined Laplacian-$k$NN |
|---|---|---|---|---|
| Prediction neighbors, $k_p$ | 1 | 0 (0%) | 6 (12%) | 3 (6%) |
| | 3 | 2 (4%) | 27 (54%) | 22 (44%) |
| | 5 | 40 (80%) | 17 (34%) | 25 (50%) |
| | 10 | 8 (16%) | 0 (0%) | 0 (0%) |
| | 20 | 0 (0%) | 0 (0%) | 0 (0%) |
| | 40 | 0 (0%) | 0 (0%) | 0 (0%) |
| Graph neighbors, $k_g$ | 10 | – | 25 (50%) | 19 (38%) |
| | 20 | | 0 (0%) | 24 (48%) |
| | 40 | | 2 (4%) | 7 (14%) |
| | 80 | | 23 (46%) | 0 (0%) |
| Laplacian weight, $\mu$ | 0.01 | – | 0 (0%) | 0 (0%) |
| | 0.1 | | 25 (50%) | 37 (74%) |
| | 1 | | 25 (50%) | 13 (26%) |
| | 10 | | 0 (0%) | 0 (0%) |

Table S3 – Connectivity properties of the final Laplacian graphs. Values are mean ± standard deviation across 50 final refitted graphs (10 folds over 5 random seeds).

| Graph property | Labeled Laplacian-*k*NN | Combined Laplacian-*k*NN |
|---|---|---|
| Graph nodes | 24,892 ± 0.4[a] | 41,079 ± 0.4 |
| Labeled nodes | 24,892 ± 0.4 | 24,892 ± 0.4 |
| Unlabeled nodes | – | 16,187 |
| Labeled-labeled edges (%) | 100 ± 0.0 | 44.6 ± 1.5% |
| Labeled-unlabeled edges (%) | – | 32.4 ± 2.9% |
| Unlabeled-unlabeled edges (%) | – | 22.9 ± 1.4% |
| Mean weighted degree[b] | 26.2 ± 20.8 | 11.9 ± 6.0 |
| Unlabeled nodes in unanchored components[c] | – | 0.57 ± 0.41% |

a. The number of labeled graph nodes corresponds to the training set in each fold (90% of the 27,658 labeled records). The remaining 10% (approximately 2,766 records) were held out for testing and excluded from graph construction.
b. Mean weighted degree is the average, across graph nodes, of the sum of incident edge weights, $d_i = \sum_j w_{ij}$, and therefore reflects the overall strength of local graph connectivity.
c. Unlabeled nodes in unanchored components denote the percentage of unlabeled nodes belonging to connected components that contain no labeled node and therefore have no path to a labeled anchored target through the graph.

Table S4 – Predictive performance for $k_{cat}$ in the adenylate kinase dataset. Performance is reported for the neural consistency and graph-Laplacian frameworks under supervised, labeled regularization, and combined labeled–unlabeled conditions across the 30 predefined ADK train/test splits. Values are mean ± standard deviation across splits. Bold values indicate the best performing condition within for each metric. Statistical significance was assessed using the paired Wilcoxon signed-rank test across the 30 splits. Asterisks indicate significant improvement relative to the supervised baseline (*p < 0.05, **p < 0.01, ***p < 0.001) whereas daggers indicate significant improvement relative to the corresponding labeled-only regularization condition (†p < 0.05, ††p < 0.01, †††p < 0.001). Significance symbols are shown in parentheses beneath the corresponding values.

| Training Condition | | RMSE | $R^2$ | PCC | $\rho$ |
|---|---|---|---|---|---|
| Neural Network | Supervised | 0.560 ± 0.052 | 0.146 ± 0.149 | 0.424 ± 0.163 | 0.360 ± 0.151 |
| | Labeled-only | 0.558 ± 0.052 (***) | 0.150 ± 0.151 (***) | 0.426 ± 0.164 (*) | **0.360 ± 0.152** |
| | Combined | **0.559 ± 0.051** (**) | **0.150 ± 0.150** (**) | **0.426 ± 0.163** (*) | 0.359 ± 0.154 |
| *k*-Nearest Neighbor | Supervised | 0.575 ± 0.058 | 0.108 ± 0.122 | 0.401 ± 0.117 | 0.311 ± 0.124 |
| | Labeled-only | 0.573 ± 0.057 | 0.116 ± 0.114 | 0.400 ± 0.127 | 0.313 ± 0.130 |
| | Combined | **0.569 ± 0.048** | **0.121 ± 0.141** | **0.404 ± 0.162** | **0.357 ± 0.133** |

Table S5 – Predictive performance for $K_m$ in the adenylate kinase dataset. Performance is reported for the neural consistency and graph-Laplacian frameworks under supervised, labeled regularization, and combined labeled–unlabeled conditions across the 30 predefined ADK train/test splits. Values are mean ± standard deviation across splits. Bold values indicate the best performing condition within for each metric. Statistical significance was assessed using the paired Wilcoxon signed-rank test across the 30 splits. Asterisks indicate significant improvement relative to the supervised baseline (*p < 0.05, **p < 0.01, ***p < 0.001) whereas daggers indicate significant improvement relative to the corresponding labeled-only regularization condition (†p < 0.05, ††p < 0.01, †††p < 0.001). Significance symbols are shown in parentheses beneath the corresponding values.

| Training Condition | | RMSE | $R^2$ | PCC | $\rho$ |
|---|---|---|---|---|---|
| Neural Network | Supervised | 0.351 ± 0.043 | −0.135 ± 0.158 | 0.121 ± 0.114 | 0.126 ± 0.110 |
| | Labeled-only | 0.350 ± 0.042 | −0.132 ± 0.154 | **0.123 ± 0.114** | **0.128 ± 0.109** |
| | Combined | **0.350 ± 0.042** (*) | **−0.130 ± 0.151** (*) | 0.121 ± 0.113 | 0.125 ± 0.110 |
| *k*-Nearest Neighbor | Supervised | 0.340 ± 0.046 | −0.069 ± 0.228 | 0.229 ± 0.140 | 0.237 ± 0.129 |
| | Labeled-only | 0.335 ± 0.042 | −0.041 ± 0.194 | 0.238 ± 0.144 | 0.260 ± 0.136 |
| | Combined | **0.326 ± 0.041** (**,†) | **0.015 ± 0.153** (**,†) | **0.280 ± 0.196** (*) | **0.308 ± 0.196** (**) |

Table S6 – Final objective functions of the training conditions used in the neural consistency regularization experiment

| Training Condition | $\lambda_L$ | $\lambda_U$ | Objective Function |
|---|---|---|---|
| Supervised | 0 | 0 | $\mathcal{L}_{\text{sup}}$ |
| Labeled consistency | 1 | 0 | $\mathcal{L}_{\text{sup}} + r(t)\mathcal{L}_{\text{L}}$ |
| Combined consistency | 1 | 1 | $\mathcal{L}_{\text{sup}} + r(t)\mathcal{L}_{\text{L}} + r(t)\mathcal{L}_{\text{U}}$ |

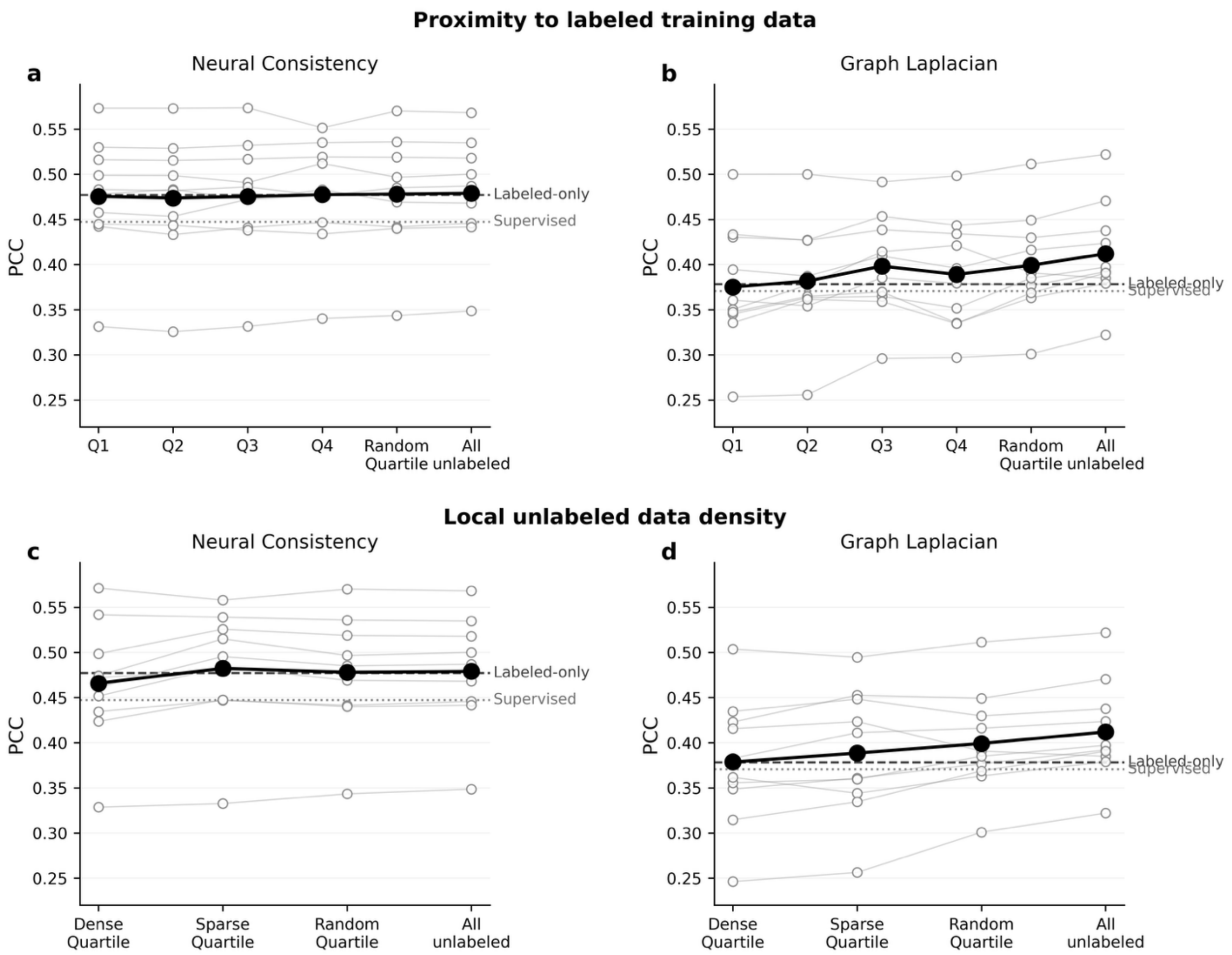


Figure S1 – Dependence of SSL performance on the feature representation space of unlabeled data using PCC as a metric. (a, b) Unlabeled observations were ranked by their mean Euclidean distance to the 20 nearest labeled training observations and divided into four approximately equal proximity quartiles, from Q1 (closest) to Q4 (farthest). Neural consistency is shown in (a) and graph Laplacian regularization in (b). (c, d) Local unlabeled data density was quantified using the mean Euclidean distance to the 20 nearest unlabeled observations. The dense quartile contained the 25% of records with the smallest mean neighbor distances whereas the sparse quartile contained the 25% with the largest distances. Neural and graph results are shown in (c) and (d), respectively. A randomly sampled quartile and the complete set of 16,187 unlabeled observations were included as reference conditions. Open gray circles represent individual fold performances after averaging across five random seeds, with thin gray lines connecting results from the same fold. Large black circles and connecting lines indicate the mean across the ten folds. Dashed horizontal lines indicate the labeled consistency or regularization condition while dotted lines indicate the corresponding supervised baseline.

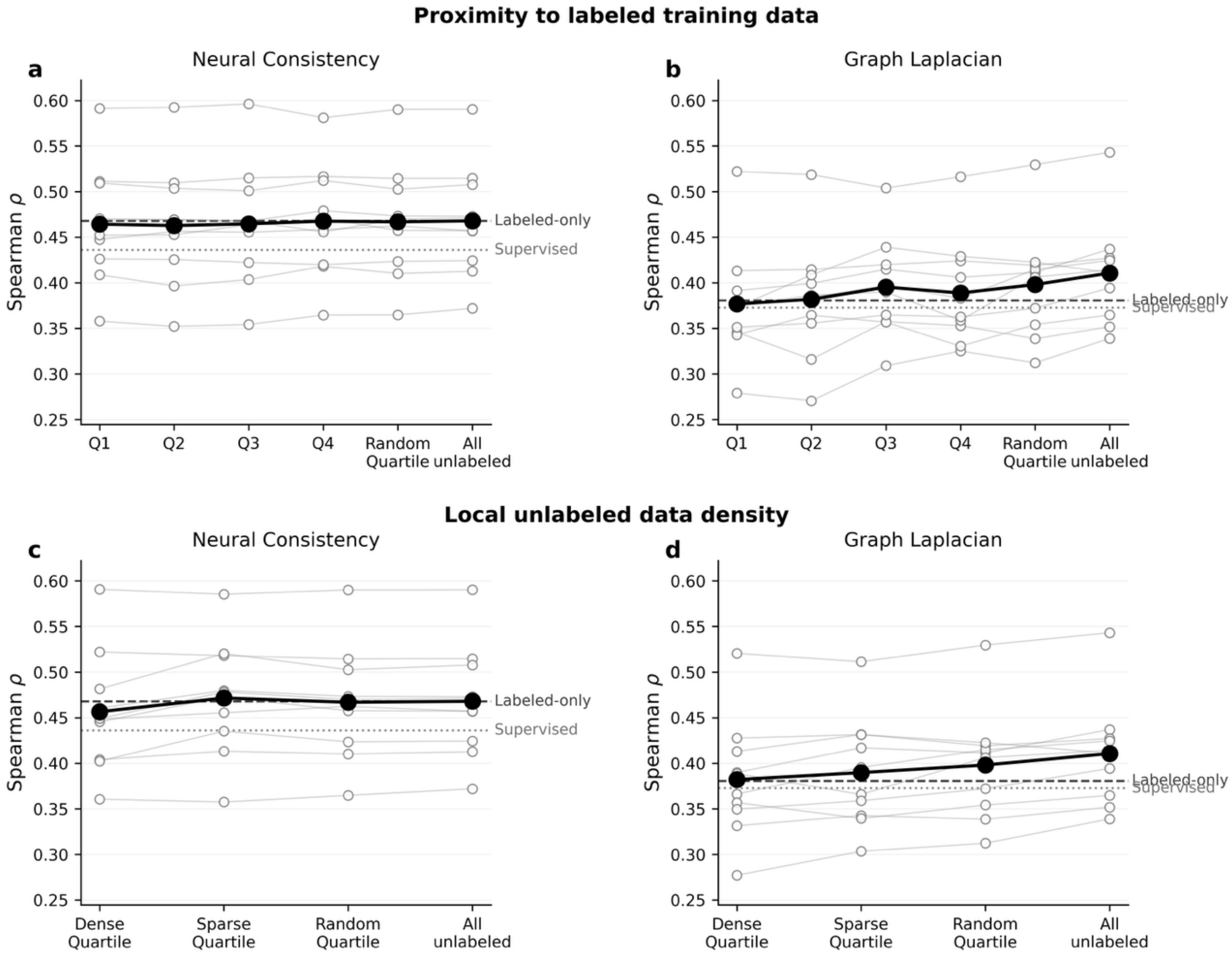


Figure S2 – Dependence of SSL performance on the feature representation space of unlabeled data using Spearman's $\rho$ as a metric. (a, b) Unlabeled observations were ranked by their mean Euclidean distance to the 20 nearest labeled training observations and divided into four approximately equal proximity quartiles, from Q1 (closest) to Q4 (farthest). Neural consistency is shown in (a) and graph Laplacian regularization in (b). (c, d) Local unlabeled data density was quantified using the mean Euclidean distance to the 20 nearest unlabeled observations. The dense quartile contained the 25% of records with the smallest mean neighbor distances whereas the sparse quartile contained the 25% with the largest distances. Neural and graph results are shown in (c) and (d), respectively. A randomly sampled quartile and the complete set of 16,187 unlabeled observations were included as reference conditions. Open gray circles represent individual fold performances after averaging across five random seeds, with thin gray lines connecting results from the same fold. Large black circles and connecting lines indicate the mean across the ten folds. Dashed horizontal lines indicate the labeled consistency or regularization condition while dotted lines indicate the corresponding supervised baseline.

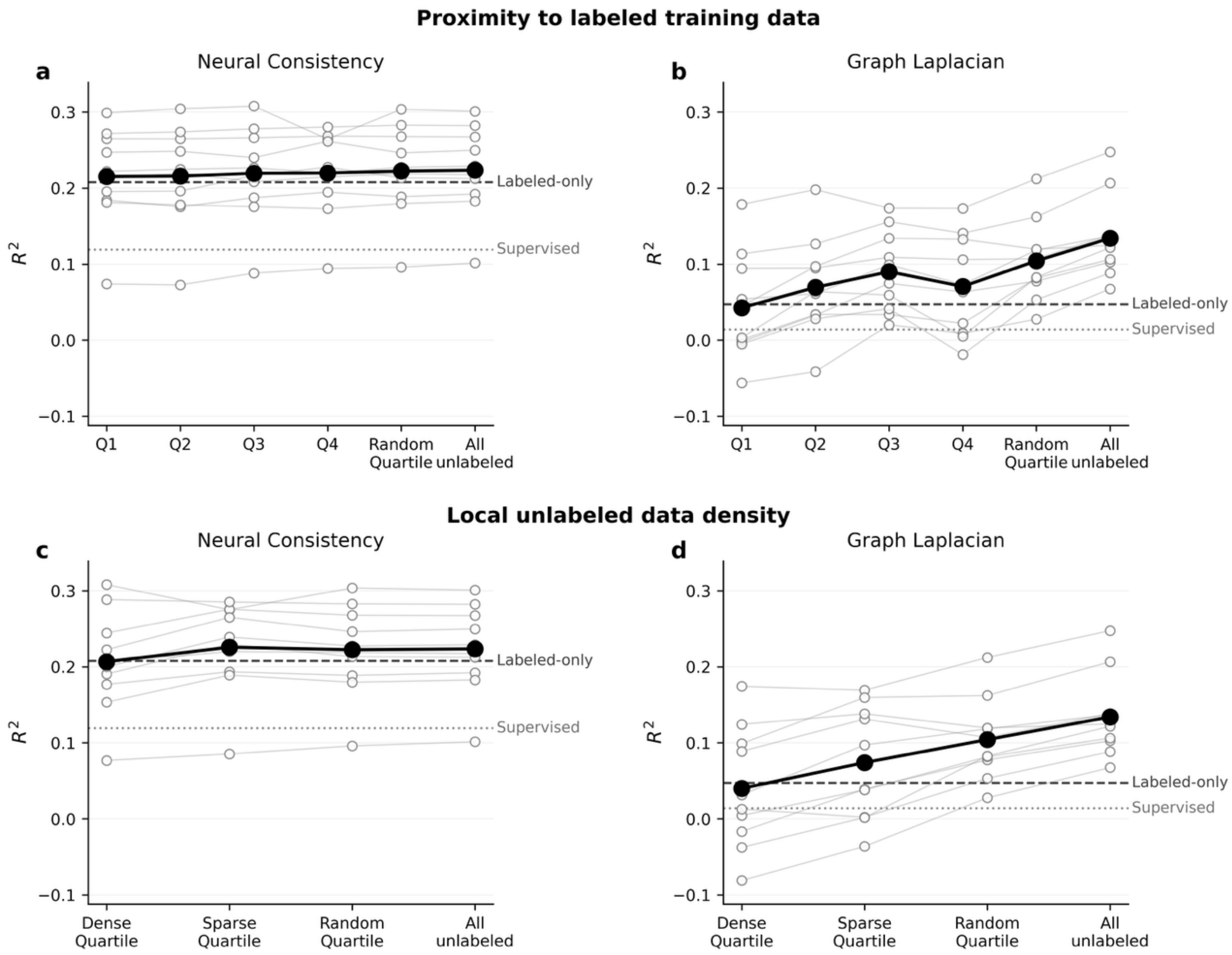


Figure S3 – Dependence of SSL performance on the feature representation space of unlabeled data using $R^2$as a metric. (a, b) Unlabeled observations were ranked by their mean Euclidean distance to the 20 nearest labeled training observations and divided into four approximately equal proximity quartiles, from Q1 (closest) to Q4 (farthest). Neural consistency is shown in (a) and graph Laplacian regularization in (b). (c, d) Local unlabeled data density was quantified using the mean Euclidean distance to the 20 nearest unlabeled observations. The dense quartile contained the 25% of records with the smallest mean neighbor distances whereas the sparse quartile contained the 25% with the largest distances. Neural and graph results are shown in (c) and (d), respectively. A randomly sampled quartile and the complete set of 16,187 unlabeled observations were included as reference conditions. Open gray circles represent individual fold performances after averaging across five random seeds, with thin gray lines connecting results from the same fold. Large black circles and connecting lines indicate the mean across the ten folds. Dashed horizontal lines indicate the labeled consistency or regularization condition while dotted lines indicate the corresponding supervised baseline.
